# Comparing the Influence of Habitat Configuration on Population Connectivity and Genetic Structure Using Congeneric Species Across Multiple Taxa

**DOI:** 10.64898/2026.09.25.754572

**Authors:** Masaki Takenaka, Seiya Okamoto, Hirohisa Suzuki, Takumi Yoshida, Gaku Ueki, Koji Tojo

**Author notes:** co-first authors (Masaki Takenaka, Seiya Okamoto). Corresponding Author: Masaki Takenaka.

## Abstract

Habitat configuration influences population connectivity and, consequently, genetic structure. River networks provide heterogeneous, hierarchically arranged environments that shift drastically from upstream to downstream. We compared congeneric species across Ephemeroptera, Plecoptera, and Trichoptera, emphasizing longitudinal replacement (upstream to downstream) and the rarely studied wet rock (hygropetric) habitats. We surveyed six rivers on the Muroto Peninsula, Japan, qualitatively sampling aquatic insects at 63 sites. Cluster analysis based on the environmental data classified surveyed sites into four clusters. We analyzed a total of 10 species from four genera that exhibit longitudinal replacement patterns within genera. Genetic analyses based on the mitochondrial cytochrome c oxidase subunit I region revealed that upstream species showed higher genetic diversity than downstream species. In contrast, species adapted to hygropetric exhibited the lowest genetic differentiation among all habitat types. A novel contribution of this study is the inclusion of hygropetric species. The surprisingly low differentiation in hygropetric species suggests high connectivity, similar to lentic species. By comparing congeneric taxa across orders within environmentally similar rivers, we reduce phylogenetic and environmental confounds, strengthening inference that habitat configuration and dispersal traits jointly shape genetic structure. These findings provide a new perspective on riverine spatial ecology and underscore the importance of microhabitat-aware comparisons for evolutionary inference.

## Introduction

Understanding the spatial distribution pattern of genetic diversity is a central and critical topic in the study of biodiversity, and also in maintaining biodiversity (Zhou et al. 2008; Hughes et al. 2013; Paz-Vinas et al. 2015; Takenaka et al. 2021). The spatial pattern of genetic structure has been reported in several studies (Paz-Vinas et al. 2015; de Araujo Barbosa et al. 2025), including patterns such as genetic structure resulting from isolation by distance (Wright 1943; Sexton et al. 2014; Takenaka and Tojo 2019), genetic structure shaped by distance along adapted habitats (Murray et al. 2019; Fusco et al. 2021; Ueki and Tojo 2023), genetic structure influenced by the spatial configuration of adapted habitats (Hof et al. 2012; Hughes et al. 2013; Takenaka et al. 2019), and genetic structure influenced by dispersal ability (Bowler and Benton 2005; Ikeda et al. 2012; Ohnishi et al. 2021). Genetic structure is determined by the complex interaction of various influences. Therefore, when multiple factors act in combination, it becomes difficult to assess which spatial distribution patterns are affecting genetic structure.

River systems are particularly suitable for investigating spatial patterns of genetic diversity because riverine organisms require freshwater habitats for all or part of their life cycle (Hughes et al. 2013; Waters et al. 2015; Takenaka et al. 2021). Therefore, river networks include highly heterogeneous environments and hierarchical patterns, and the environmental characteristics of rivers drastically transition from upstream to downstream areas (Fig. 1; Campbell and McIntosh 2013; Miyazono and Taylor 2013). Also, the physical attributes of the landscape (topography, soil composition, etc.) directly influence the distributional patterns and population structures of species (Kindlmann and Burel 2008; Alp et al. 2012; Hughes et al. 2013).

**Figure 1.**
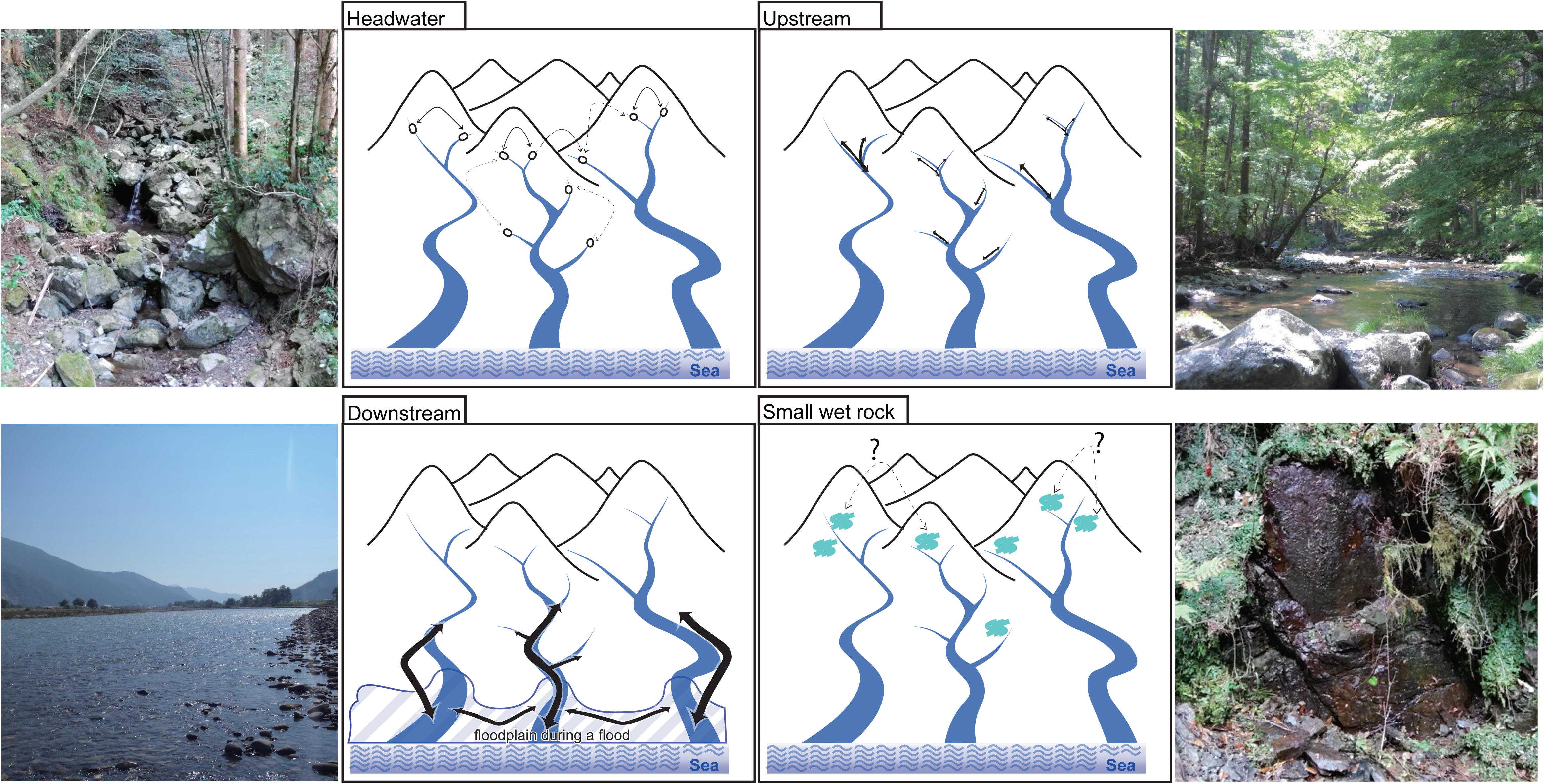
Comparison of dispersal schemes of aquatic insects adapted to each environment in the river.

Regarding organisms inhabiting river, each species inhabits different environments along the river flow, and even within the same genus, species inhabit different environments, allowing their coexistence within the same river (Fig. 1; Okamoto and Tojo 2021; Okamoto et al. 2022). Thus, riverine organisms provide an ideal system for examining the relationship between habitat spatial configuration and genetic structure. By comparing genetic structures among multiple taxa, Hughes et al. (2013) tries to make broad generalizations about population connectivity based on species’ life history and the structure of the aquatic habitat. Species adapted to upstream environments often inhabit scattered and isolated habitats, and this patchy distribution increases the potential for genetic differentiation among populations when compared with species adapted to downstream environments (Fig. 1; Hughes et al. 2013). However, many studies are limited to comparisons within specific taxonomic groups or comparisons between different taxonomic groups. Furthermore, species inhabiting headwater areas or small wet rock environments, known as hygropetric environments, where thin films of water flow over rocky substrates (Fig. 1, S1; Wantzen and Junk 2009), are rarely targeted. Wet rock environments differ substantially from flowing water habitats, including headwater streams, because their available habitat is smaller and more isolated. Therefore, these species are expected to exhibit greater genetic differentiation among populations compared to upstream or headwater species. However, such patterns have not been reported so far.

The genetic structure of aquatic organisms is shaped by a complex interplay of factors, including habitat connectivity and species’ dispersal capacity (Ikeda et al. 2012; Hughes et al. 2013; Hjalmarsson et al. 2015; Takenaka and Tojo 2019; Waters et al. 2020). When dispersal capacity is high, frequent gene flow among populations may occur, resulting in more homogeneous genetic structures (Sproul et al. 2014; Phillipsen et al. 2015). While direct evaluation of dispersal ability among species remains challenging (Ohba et al. 2025), the scale of realized gene flow can be assessed through genetic data. Even in species with high flight capability, dispersal may be ineffective unless individuals can successfully establish in new habitats; therefore, estimating genetic dispersal capacity is more informative than evaluating flight ability alone.

According to the models of dispersal capacity and population structure (detail in Finn et al. 2007, Hughes et al. 2009, 2013), species adapted to headwater environments generally correspond to the Headwater Model (HM) (Takenaka et al. 2019; Lancaster et al. 2024). The HM suggests that, despite the patchy and spatially restricted nature of headwater habitats, genetic connectivity may be maintained among some populations through dispersal between catchments, a pattern that has been documented for several taxa (Finn et al. 2007; Hughes et al. 2009). However, because wet rock habitat is often more spatially isolated than headwater streams, we expect these species to exhibit genetic patterns consistent with the Death Valley Model (DVM), which assumes physically isolated environments with little surface-water connectivity, resulting in strong genetic structure among sites. In contrast, downstream species are predicted to approach the Panmixia Model (PAN). If dispersal among river is partially restricted, the Stream Hierarchy Model (SHM) may also apply. Nevertheless, for flying aquatic insects, their relatively high dispersal capacity should result in genetic structures closer to PAN compared with species adapted to more isolated habitats such as upstream headwaters or the wet rock habitat. Species inhabiting intermediate environments, such as mid- and upper-stream, are expected to conform to the SHM.

To evaluate our hypotheses, it is important to control for other variables. In this study, we focused on closely related species, as they share many common factors. However, even related species may have followed different evolutionary paths. To comprehensively assess the spatial effects of habitat on genetic structure, we compared congeneric species across multiple taxonomic groups. Therefore, our study examines multiple congeneric species within a single peninsula. This design enables us to investigate the relationship between species-specific habitat use and population connectivity through genetic structure, under relatively similar environmental conditions. Although, in addition, previous studies have examined the effects of habitat fragmentation and human impacts on genetic structure (e.g., Monaghan et al. 2002; Davis et al. 2018), the present study focuses on rivers with a high degree of naturalness that flow relatively simply from mountainous headwaters to the sea. This allows us to discuss the relationship between the river’s intrinsic spatial configuration and genetic patterns.

Our aim was to test whether the spatial distribution of suitable habitats influences genetic structure by comparing congeneric species inhabiting different riverine environments. Specifically, we hypothesized that upstream species would exhibit greater genetic differentiation than their downstream congeners within the six rivers of the Muroto Peninsula. In addition, regarding dispersal capacity and population structure according to Hughes et al. (2009), upstream species within each genus such as *Eph. japonica*, *Epe. nipponicus*, *C. japonica*, and *Do. commata*, as well as headwater species such as *Di. tipuliformis* and *Do. japonica*, correspond to the HM. Downstream species such as *Eph. strigata*, *Epe. ikanonis*, *I. japonica*, and *S. marmorata* are predicted to fit the PAN. Wet rock species such as *Do. angustata* and *C. kawasawai* are expected to show tendencies of the DVM, because their movement among sites is considered to be restricted. Additionally, *Epe. curvatulus* is predicted to follow the SHM (Table 1).

**Table 1.**
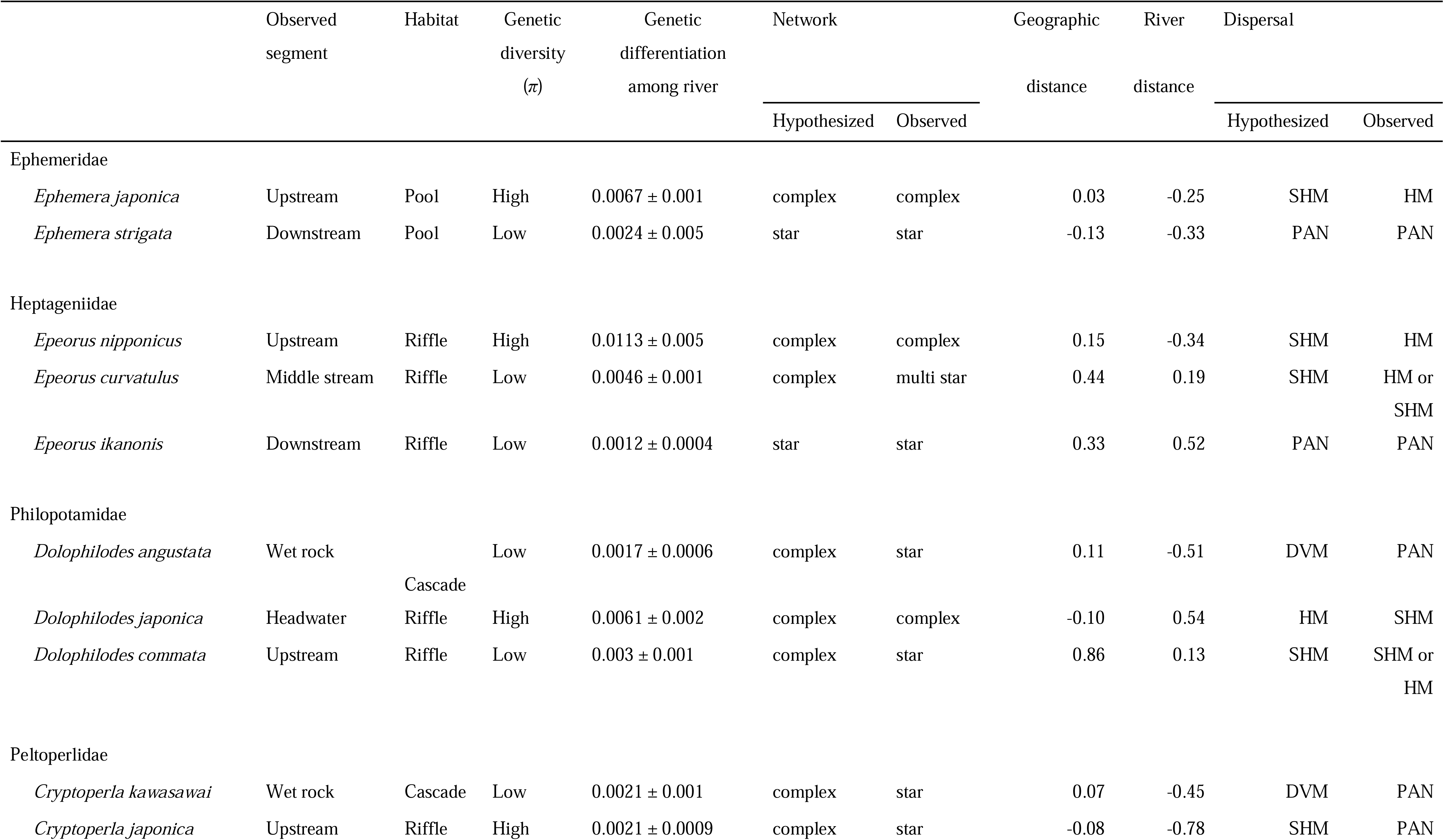
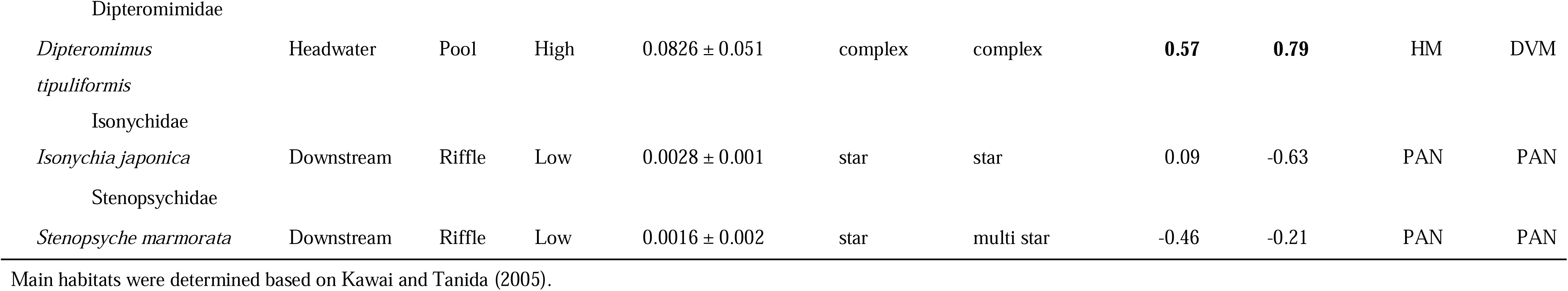
Hypotheses and results on dispersal capacity and population structure.

**Table 2.**

| Species | N | H | Hd | Pi |
| --- | --- | --- | --- | --- |
| *Epeorus nipponicus* | 23 | 14 | 0.93 | 0.00927 |
| *Epeorus curvatulus* | 29 | 22 | 0.96 | 0.00449 |
| *Epeorus ikanonis* | 16 | 4 | 0.59 | 0.00144 |
| *Ephemera japonica* | 31 | 18 | 0.94 | 0.00653 |
| *Ephemera strigata* | 27 | 8 | 0.46 | 0.00127 |
| *Dolophilodes angustata* | 33 | 6 | 0.67 | 0.00131 |
| *Dolophilodes japonica* | 24 | 18 | 0.98 | 0.00586 |
| *Dolophilodes commata* | 17 | 11 | 0.94 | 0.00319 |
| *Cryptoprla kawasawai* | 25 | 4 | 0.30 | 0.00074 |
| *Cryptoprla japonica* | 29 | 15 | 0.77 | 0.00217 |
| *Dipteromimus tipuliformis* | 27 | 17 | 0.957 | 0.06873 |
| *Isonychia japonica* | 18 | 11 | 0.882 | 0.00301 |
| *Stenopsyche marmorata* | 23 | 3 | 0.372 | 0.00201 |

## Methods

### Study Design and sampling

Regarding study sites, we focused on a single peninsula and selected six rivers on the peninsula that are relatively similar in environmental characteristics. In January and June 2022, we investigated six rivers in Muroto Peninsula (6–13 sampling sites per river; Table S1; Fig. 2). In all rivers, the number of sampling sites differed among rivers in order to collect at least five individuals per target species per river examined in this study. In addition, when some species belonging to taxonomic groups that are difficult to identify were not collected during the January survey, additional surveys were conducted in June. Within a river, in order to comprehensively cover various environments, surveys were conducted not only from the upstream to downstream areas but also in headwater and small wet rock environments (Fig. S1). Qualitative sampling for aquatic insects was conducted at 63 sites. In this study, headwaters were defined as small tributaries flowing into upstream that are not mapped as rivers on 1:25,000 topographic maps of the Geospatial Information Authority of Japan. Small wet rock environments were defined as habitats where thin films of water flow over rocky substrates. River environments, including upstream and downstream, were classified based on clusters derived from environmental data, and are generally treated as relative concepts along a longitudinal gradient from the river mouth.

**Figure 2.**
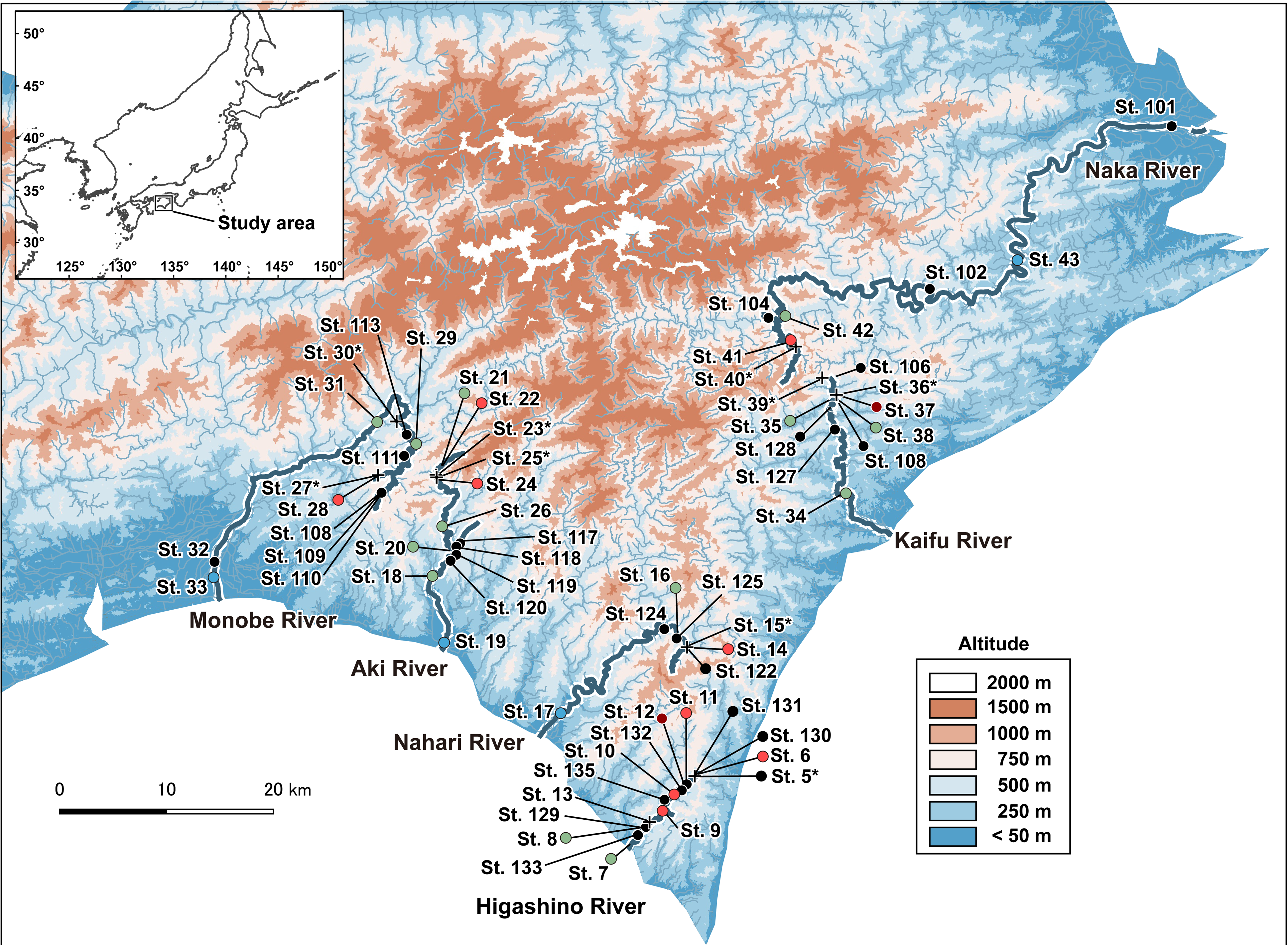
The map shows the six rivers that were the focus of this study: the Higashino River, the Nahari River, the Aki River, the Monobe River, the Naka River, and the Kaifu River, as well as the surveyed localities. The map was created using QGIS 2.1, with elevation differences illustrated through color gradation on Muroto Peninsula. The color of each circle represents the results of figure 3. The color of each circle represents the results shown in Figure 3 (Black circles were not measured environments). Small wet rock environments are indicated by crosses.

Regarding the target species, we focused on ten species across four genera [*Epeorus*, *Ephemera* (Ephemeroptera), *Cryptoperla* (Plecoptera)*, Dolophilodes* (Trichoptera)] that exhibit species replacement from upstream areas to downstream areas within the same genus, and *Dipteromimus tipuliformis* (Ephemeroptera), which is distributed in the headwater region, as well as *Isonychia japonica* (Ephemeroptera) and *Stenopsyche marmorata* (Trichoptera), which are distributed relatively downstream.

For the genus *Ephemera*, *Ephemera japonica* inhabits areas further upstream compared to *Ephemera strigata* (Okamoto and Tojo 2021). For the genus *Epeorus*, *Epeorus nipponicus* inhabits areas further upstream than *Epeorus curvatulus* (Ogitani and Nakamura 2008), while *Epeorus ikanonis*, on the other hand, tends to inhabit relatively downstream areas.

For *Dolophilodes*, we used *Dolophilodes angustata*, *Dolophilodes japonica*, and *Dolophilodes commata*(Kuhara 2005; Kuhara 2018). *Dolophilodes angustata* inhabits small wet rock environments (Kuhara 2018), and *Do. japonica* is found further upstream compared to *Do. commata*. For *Cryptoperla*, we used *Cryptoperla kawasawai*, which inhabits small wet rock environments, and *Cryptoperla japonica*, which is found in upstream areas.

In flowing-water environments (excluding wet rock environments), all taxa were collected using a surver net (net width of 250 mm, and mesh size of 0.493 mm: NGG38). In wet rock environments, specimens were collected by hand using forceps or by gently picking them up with fingers. Species identification was conducted following Kawai and Tanida (2018).

### Environmental factors measured at each study site

To evaluate the environmental factors at 39 study (surveyed locations in January, 2022) sites of 63 sites, the following factors were obtained using field surveys and geographic information system (GIS): the underlying elevation data (Altitude) were obtained from a 5 m or 10 m digital elevation model (DEM) provided by the Geospatial Information Authority of Japan using GIS. Riverbed slope degree was calculated using GIS mapping based on the distance between points at which there was a 10 m elevation difference, with each study site set as the base point. This analysis used the 5 m or 10 m DEM provided by the Geospatial Information Authority of Japan. Channel width was measured using a measuring tape. For sites St. 17, St. 19, St. 32, St. 33, St. 34, and St. 43 where measurement of width was difficult because the channel was relatively large and water depth was deep, the widths were measured using GIS. Canopy openness was measured using CanopOn2 software (http://takenaka-akio.org/etc/canopon2/) and a digital hemispherical picture taken with a fisheye lens (EX-FR200, CASIO, Tokyo, Japan) at the center of each study site. In some cases, the periphery of the hemispherical pictures included distant artificial objects in the background. However, the objects did not affect openness the over the river channel. Thus, the canopy openness was calculated, after excluding 10% of the data corresponding to the periphery/edges of all of photographs to eliminate such “noise”. Substrate coarseness was assessed the five transect lines along the river and five transect lines across the river, and the grain size was measured at these intersections, resulting in 25 measurement points per site (see Okamoto et al. 2022). Grain size was measured at each measurement point and each substrate sample was classified as follows: sand and silt (particle size < 2 mm), gravel (2–16 mm), pebble (17–64 mm), cobble (65–256 mm), and boulder (> 256 mm). Subsequently, we calculated average grain size for the 25 measurement points using the following numeric classification protocol (Bain et al. 1985; Saito and Tojo 2016): sand or silt = 1, gravel = 2, pebble = 3, cobble = 4, boulder = 5.

### Clustering analysis

To compare the environmental factors among 39 study sites, hierarchical cluster analysis was performed based on Euclidean distances, along with the Ward method. The software R ver. 4.2.1 was used for this analysis (R Core Team 2020). At elevations and along the longitudinal course of the river, riverbed slope, river width, substrate coarseness, and canopy openness changed. Notably, the canopy openness was strongly negatively correlated to elevation (r = −0.78). Because the targeted rivers varied in drainage basin size and characteristics, a preliminary classification was conducted based on environmental variables (riverbed slope degree, channel width, and substrate coarseness, excluding highly correlated variables, r > 0.7) using cluster analyses.

### DNA data

In the laboratory, total genomic DNA was extracted from ethanol-preserved tissues of specimens and purified using a DNeasy Blood & Tissue Kit (QIAGEN, Hilden) or GenCheck® DNA Extraction Reagent (FASMAC, Kanagawa), according to the manufacturer’s instructions. Each total genomic DNA was used to amplify DNA fragments [the mitochondrial DNA (mtDNA) cytochrome c oxidase subunit I (COI)] by polymerase chain reaction (PCR) using a set of primers LCO1490 (5’– GGTCAA CAA ATC ATA AAG ATA TTG G –3’), HCO2198 (5’– TAA ACT TCA GGG TGA CCA AAA AAT CA–3’) and HCOoutout (5’- GTA AAT ATA TGR TGD GCT C –3’) (Folmer et al. 1994; Prendini et al. 2005). The following PCR protocol was used: 92℃ for 2 min; 35× (94℃ for 1 min, 48℃ or 50℃ for 30 sec, 72℃ for 1min; 72℃ for 3min). PCR products were purified using ExoSAP-IT or illustra ExoProStar (GE Healthcare, Buckinghamshire). Purified DNA fragments were sequenced using a BigDye Terminator v1.1 Cycle Sequencing Kit (Applied Biosystems, California) on a DNA Sequencer (ABI 3130 or 3130xl DNA Analyzer; Perkin Elmer/Applied Biosystems, California).

All sequence data were submitted to the DNA databank of Japan (DDBJ database; Table S2). Sequence alignment and editing were performed for each gene separately using “auto-strategy” of MAFFT ver. 7 (Katoh and Standley 2013) and CLC Workbench software (CLC bio, Aarhus, Denmark). The alignments were determined for unique haplotypes and genotypes using the software DnaSP ver. 4.0 (Rozas et al. 2003) prior to subsequent analysis. Finally, the sequences of all species were trimmed to 633 bp and used in downstream analyses.

### Genetic structure

Haplotype networks were constructed to visualise genetic relationships among individuals and/or populations and to assess whether genetic structure corresponded to the spatial distribution of habitats occupied by each species. Haplotype networks were constructed by the program TCS Network (Clement et al. 2000) based on the mtDNA COI region (633 bp) using PopART (Leigh et al. 2015). The networks were colour-coded by habitat type based on environmental clustering of the sampling sites (Sampling site groups), with habitats classified as headwater, upstream, downstream, and middle stream, defined as environments between upstream and downstream. For the degree of genetic diversity (haplotype diversity: *Hd*, and nucleotide diversity: *Pi*), intraspecific genetic diversity was analysed for each species and each sites using software DnaSP ver. 4.0 (Rozas et al. 2003). The resulting nucleotide diversity values for each study sites were visualized using box plots. Genetic differentiation among river in each species were calculated as genetic distance (*p*-distance) using MEGA ver. 7 (Kumar et al. 2016).

### Data Analysis

To identify the nucleotide diversity in each species, we used a generalized liner model (GLM) with Gaussian distribution. We ran a GLM with nucleotide diversity (numerical) for each species as the response variable and species as explanatory factors (categorical). The statistical analyses were conducted in R 4.0.1 (R Core Team 2020). Genetic distances among specimens were calculated using *p*-distance, and the mean genetic distance was subsequently calculated for each site. We evaluated the correlation between genetic distance and two types of distance among population using a Mantel test based on following three distance matrices: 1) pairwise genetic distance (mean p-distance) among populations, 2) pairwise Euclidean distance (geographic distance), 3) pairwise river distance. The significance of mantel *r* values was tested using 10,000 permutations performed in the R package ‘vegan’ for R. Genetic distance (*p*-distance) and Euclidean distance were calculated by AIS ver. 1.0 (Alleles In Space, Miller 2005) and river distance was measured by the R package ‘riverdist’ ver. v.0.15.0 for R (Tyers 2017).

## RESULTS

### Environments of each study site

This cluster analysis identified 4 groups based on the environmental factors (Table S3; Fig. 3, S2). In particular, Group 1 were characterized by steep riverbed slope degrees and narrow channel widths. Group 2 were characterized by narrow channel widths and substrate dominated by cobbles and boulders. Group 3 were characterized by milder riverbed slope degrees and substrates dominated by cobbles and pebbles. The sites in each targeted river were included in either Group 2 or Group 3. Group 4 were characterized by wide river channels and substrates dominated by gravel and pebbles.

**Figure 3.**
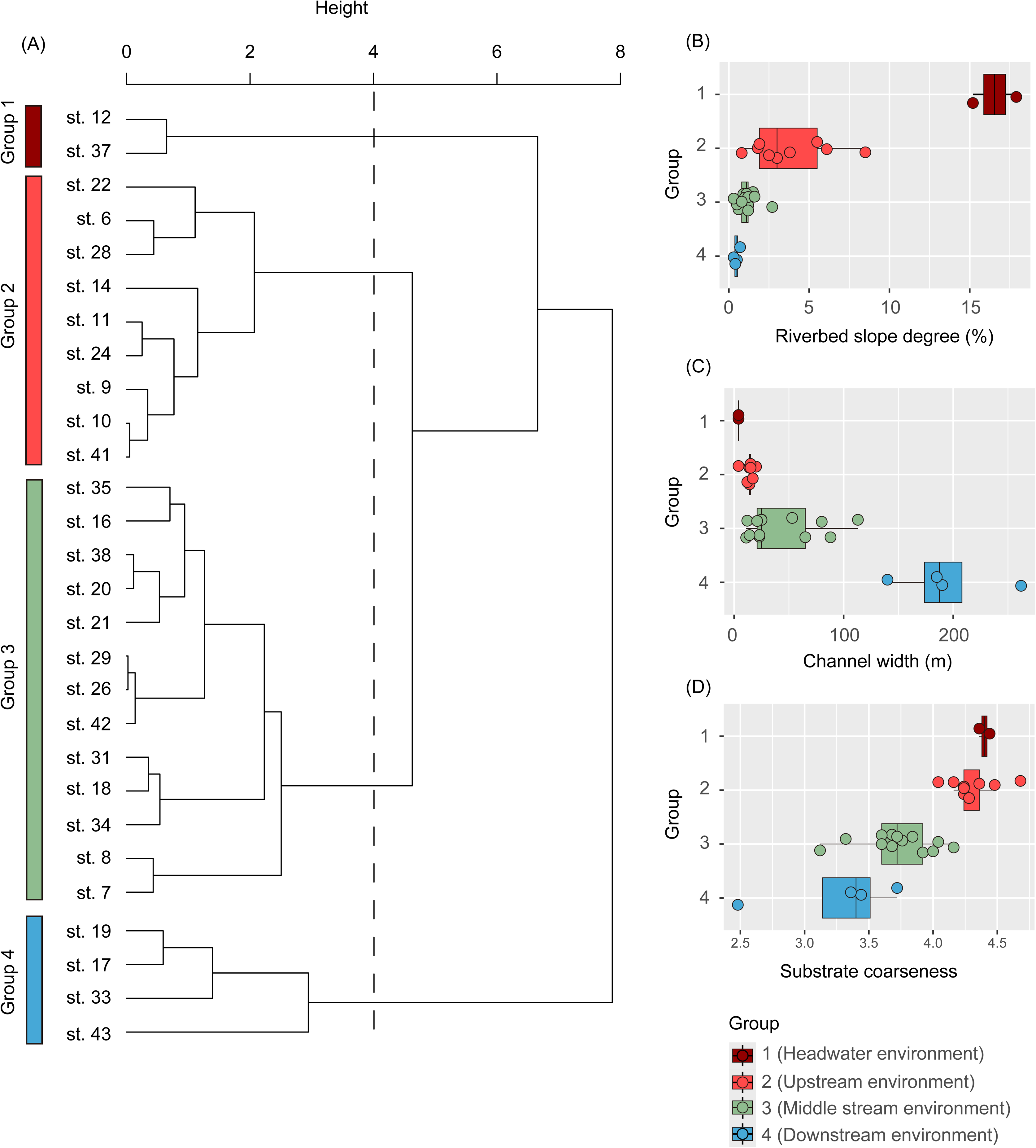
Hierarchal clustering based on three environmental factors (i.e., Riverbed slope degree, River width, Substrate coarseness) (A); the range of riverbed slope degree in each group (B); the range of channel width in each group (C); the range of substrate coarseness in each group (D)

### Genetic structure

Regarding the haplotype network, genetic structure of *Epe. nipponicus* and *Eph. japonica*, which inhabit relatively upstream areas, identified many intermediate haplotypes and detected the existence of hypothetical haplotypes that were not sampled nor known to be extinct (Fig. 4). On the other hand, the genetic structure of *Epe. ikanonis* and *Eph. strigata*, which inhabited relatively downstream areas, resulted in a simple network structure, and the genetic structure of *Epe. curvatulus*, which inhabits intermediately between these two species, was also intermediate between those of the two species. Also, nucleotide diversity and genetic distance among river, which indicate the degree of genetic differentiation, revealed that upstream species consistently showed higher genetic diversity than downstream species across all genera examined (*Ephemera*, *Epeorus*, *Cryptoperla*, and *Dolophilodes*) (Table 1, 2).

**Figure 4.**
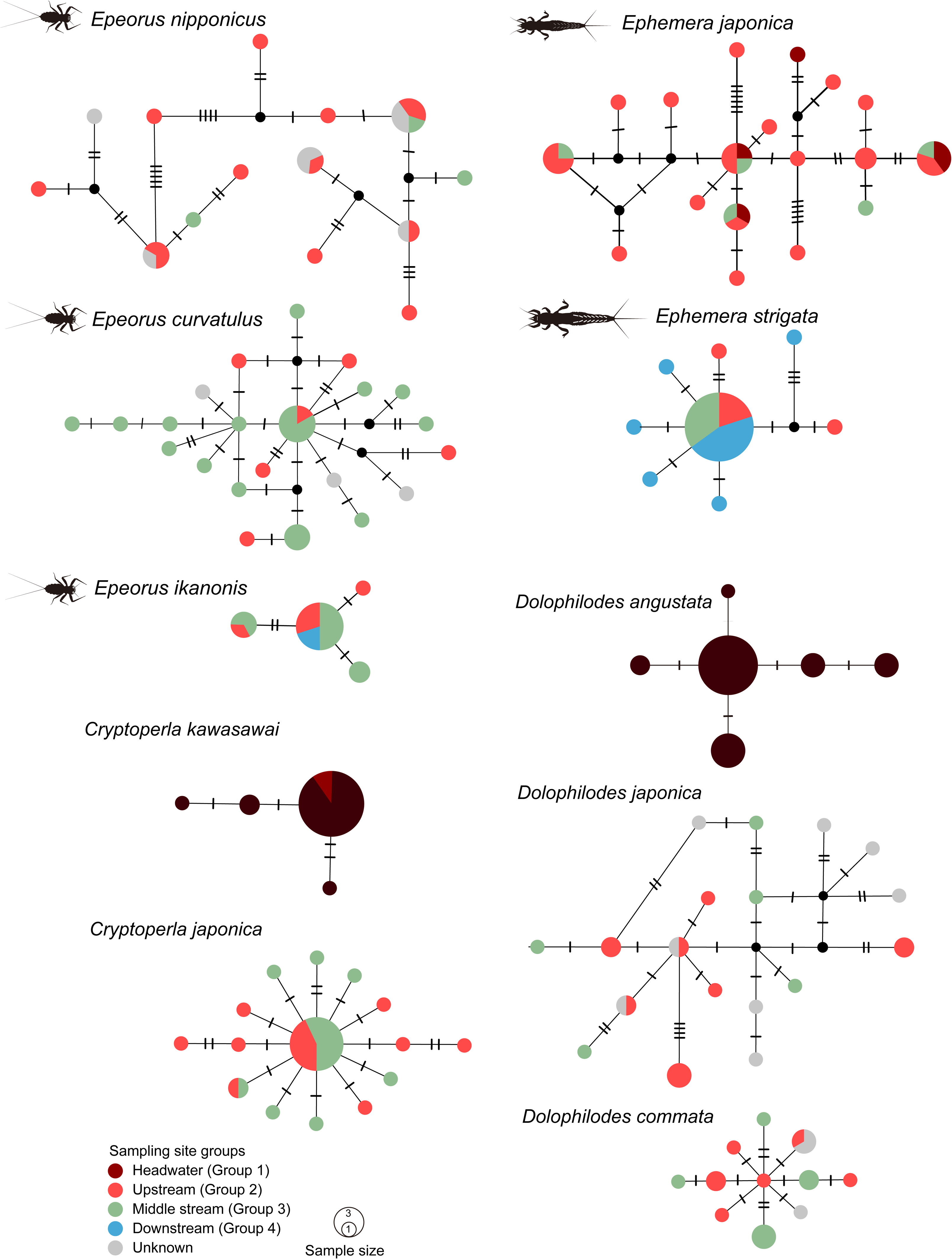
Haplotype networks based on the mtDNA COI region (701 bp) for *Epeorus* mayflies, including *Epeorus nipponicus*, *Epeorus curvatulus*, *Epeorus ikanonis*, and for *Ephemera* mayflies, including *Ephemera japonica* and *Ephemera strigata*. Haplotype networks based on the mtDNA COI region (743 bp) for *Dolophilodes* caddisflies, including the wet rock species *Dolophilodes angustata*, the headwater species *Dolophilodes japonica*, and the upstream species *Dolophilodes commata*, and for *Cryptoperla* stoneflies, including the wet rock species *Cryptoperla kawasawai* and the headwater species *Cryptoperla japonica.* The color of each circle correspond to the results of hierarchal clustering in figure 3 as follows: group 1 is headwater environments and group 2 is upstream environments, group 3 is middle stream environments, group 4 is downstream environments, and the wet rock and unknown.

To compare the genetic structure between species adapted to headwaters or upstream regions and wet rock environments within genera, we used *Dolophilodes* and *Cryptoperla* as species inhabiting these environments (Fig. 4). In the haplotype network, the genetic structure of *Do. japonica* and *C. japonica*, which inhabit headwater and upstream regions, showed many haplotypes that were relatively genetically differentiated. In contrast, the genetic structure of *Do. commata*, which inhabits areas relatively downstream, compared to *Do. japonica*, showed relatively low genetic differentiation and genetic diversity (Table 1, 2). The genetic structure of *Do. angustata* and *C. kawasawai*, which inhabit wet rock environments, showed the lowest levels of genetic differentiation.

The genetic structure of *I. japonica* and *S. marmorata*, which inhabit relatively downstream areas, showed relatively low genetic differentiation (Table 1; Fig. 5, S3). The genetic structure of *Di. tipuliformis*, which inhabits headwaters, identified many intermediate haplotypes and revealed genetic differentiation among those haplotypes (Fig. S3).

**Figure 5.**
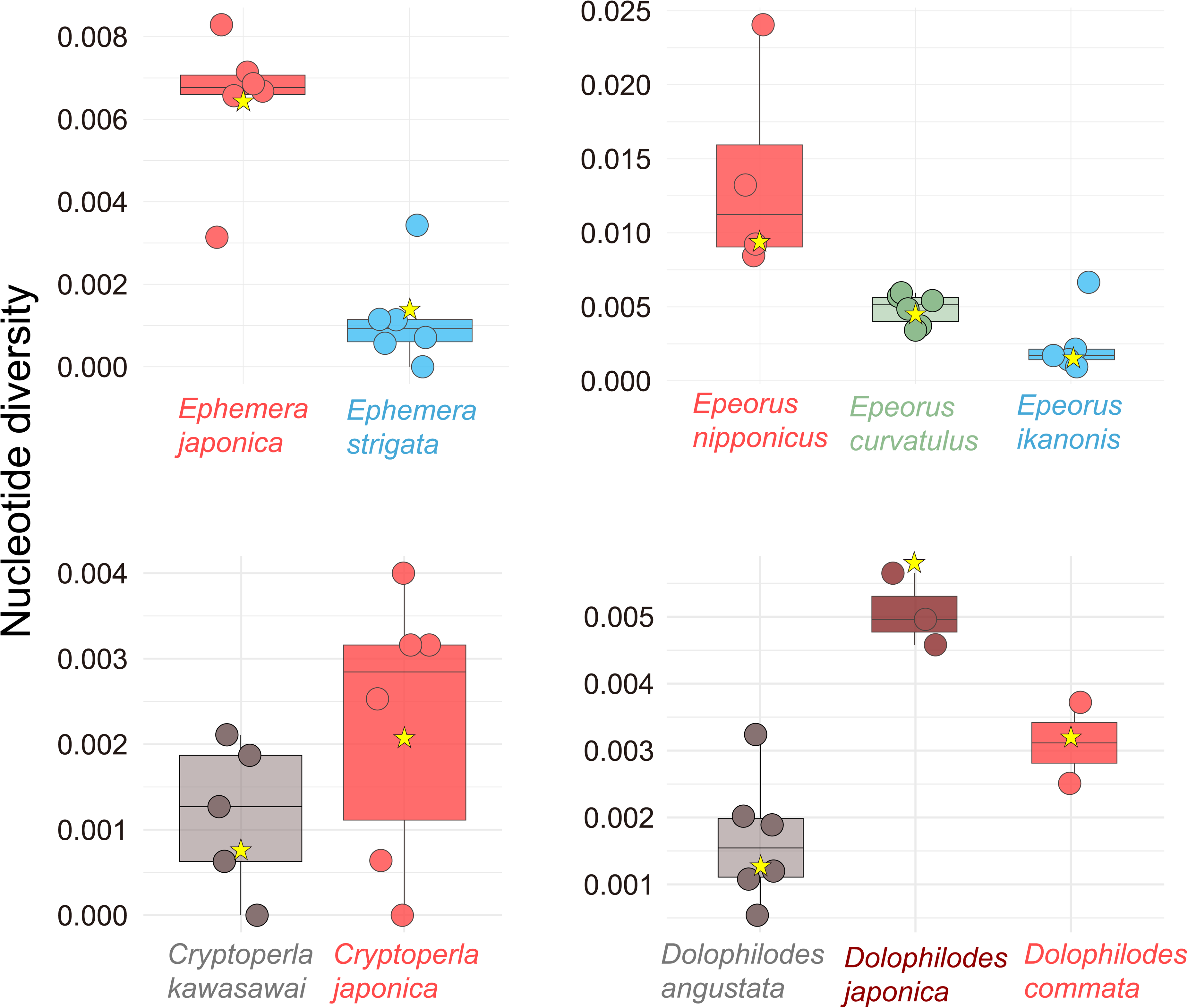
Boxplots show nucleotide diversity for each species. Yellow stars indicate the mean value for each species.

The correlation between genetic distance (*p*-distance) and geographic distance was examined (Table 1). Geographic distance was measured as both straight-line distance and river line distance calculated along the river course. The results of the estimated dispersal capacity and population structure, according to the models of Hughes et al. (2009) based on genetic structure, genetic distances, and genetic diversity, are shown in Table 1.

## Discussion

River environments change from upstream to downstream (Vannote et al. 1980; Doretto et al. 2020). The species inhabiting each environment, such as upstream and downstream, also change accordingly. The location and spatial characteristics of these habitats influence the degree and direction of gene flow between populations, which is important for understanding how the genetic structure of a species is formed. In this study, we were not able to quantitatively identify which species were present at each site. However, by comparing the environments of the survey sites and the species found there, and by examining the genetic structures of species living in the different environments, we can gain a better understanding of species distribution patterns. We analyzed the genetic structure of 13 aquatic insect species collected from several rivers that encompass a variety of environmental conditions, including small wet rock habitats differing in size and characteristics. An important advantage of this study is that it targets closely related congeneric species from three major aquatic insect orders (Ephemeroptera, Plecoptera, and Trichoptera) that inhabit different parts of the river continuum. Unlike many previous studies that claim to conduct comparative phylogeography but often compare species with substantially different evolutionary backgrounds, our approach enables high-level comparisons by focusing on ecologically and phylogenetically comparable species. This allows for a more robust evaluation of how habitat configuration influences genetic structure and population connectivity.

### Spatial Patterns of Riverine Habitats and Genetic Structure

In this study, we focused on the genera *Ephemera* and *Epeorus*, which exhibit species replacement from upstream areas to downstream areas. As supported by previous studies (Finn et al. 2007; Alp et al. 2012; Hughes et al. 2013), upstream species exhibited more complex genetic structures, while downstream species tended to show less genetic differentiation. As a spatial pattern, upstream habitats tend to be scattered and isolated, which limits gene flow between populations. Also, population sizes of species inhabiting upstream habitats are relatively smaller. Therefore, there is a higher potential for genetic differentiation between different populations compared to populations adapted to downstream environments (Hughes et al. 2013; Takenaka and Tojo 2019). Previous studies have shown that upstream or headwater species exhibit genetic differentiation among geographic regions (Yoshikawa et al. 2008; Takenaka and Tojo 2019; Mikami et al. 2023; Waters et al. 2024).

The *Di. tipuliformis*, which has adapted to headwater areas at the family level, a study conducted in the same region, the Muroto Peninsula, revealed that their genetic differentiation is significantly greater than that of the upstream species of *Ephemera* and *Epeorus* mayflies. Dipteromimid mayflies are suited to distribution in more restricted habitats (smaller scale) than the upstream species of them, which likely results in greater isolation among populations. This trend is consistent with findings from previous studies conducted in other regions (Takenaka et al. 2019). Furthermore, the genetic structures of *I. japonica* and *S. marmorata*, which are adapted to more downstream habitats, exhibit a ‘star’-like network structure, indicating high connectivity among populations. These species appear to form a single population, not only across the Muroto Peninsula, but also over a broader geographic range. Thus, this study demonstrates how spatial patterns of habitat preference can lead to differences in genetic structure.

In the genus *Ephemera*, a higher degree of genetic diversity and a more complex haplotype network were observed in *Eph. japonica* compared with *Eph. strigata*. The genetic distance of *Eph. japonica* showed a slight positive correlation with geographic distance, whereas river distance displayed a negative correlation. These patterns suggest that its dispersal capacity likely follows the HM. Because this species inhabits upstream regions and also occurs in headwaters, this interpretation is consistent with ecological expectations (Okamoto et al., 2022a, b). In contrast, low genetic diversity and simple network structure in *Eph. strigata*, indicating the PAN. In genus *Epeorus*, greater genetic diversity and a more complex network were also found in *Epe. nipponicus* in comparison with the other two species. A positive correlation was detected only for geographic distance, suggesting a high likelihood of the HM. Low genetic diversity but relatively high genetic differentiation among rivers were characteristic of *Epe. curvatulus*, and its haplotype network was complex. These results support the SHM. However, because the correlation with geographic distance was stronger than that with river distance, the possibility of the HM cannot be excluded. Low genetic distance and low genetic diversity were observed in *Epe. ikanonis*. Although both geographic and river distances showed positive correlations with genetic distance, the low overall diversity suggests that the PAN may be more appropriate. At the family level, the headwater specialist *Di. tipuliformis* inhabits only headwater environments. Although its dispersal capacity was initially predicted to fit the HM, strong genetic differentiation among sites indicates the DVM. In contrast, *I. japonica* and *S. marmorata*, which are adapted to relatively downstream habitats, showed low genetic differentiation and wide dispersal ranges, consistent with the PAN. A comparison across two genera and five species, as well as across three genera, revealed that species inhabiting upstream regions tended to show stronger genetic differentiation and were more likely to follow the HM. The stronger correlation between genetic distance and straight line geographic distance, rather than river distance, suggests frequent cross watershed movement in upstream species. Such movements across watershed divides have been directly observed in caddisflies in upstream environments (Finn et al. 2011; Lancaster et al. 2024). Downstream species generally showed lower genetic differentiation and simple genetic structures, indicating a tendency toward the PAN. Occasional correlations with geographic distance were detected, but these likely reflect effects due to minimal genetic differentiation. Previous studies reported stronger genetic differentiation in upstream species (Baggiano et al. 2011; Polato et al. 2017). In addition, species highly adapted to extreme headwater environments tended to follow the DVM. Although only one such example was found in this study, similar patterns have been reported for high altitude mountain and subalpine species (Finn and Adler 2006; Hughes et al. 2009; Pauls et al. 2006).

### Wet rock environments

Wet rock habitats, known as hygropetric environments, differ substantially from the flowing water environments of rivers (Wantzen and Junk 2009). The environmental parameters measured in this study could not be assessed in wet rock habitats (Wantzen and Junk 2009). The populations inhabiting wet rock habitats are qualitatively different. While headwater regions, though small, exist as continuous areas, wet rock habitats are spatially discrete and exist as isolated patches within the river system. This is because the downstream areas of wet rock habitats are part of the typical flowing environments, such as headwaters, which differ environmentally. Therefore, when designing this study, we hypothesized that populations inhabiting wet rock habitats would exhibit strong genetic differentiation, similar to or greater than that observed in upstream species.

Unexpectedly, species adapted to wet rock habitats showed little genetic differentiation. This result appears to align with the patterns observed in species inhabiting lentic (still) water (Arribas et al. 2012). The differences between lotic (running) water and lentic (still) water habitats are expected to influence gene flow, genetic drift, and other interpopulation-level processes due to differences in habitat connectivity (Arribas et al. 2012; Hof et al. 2012; Waters et al. 2015; Takenaka et al. 2021). Species inhabiting patchily distributed lentic water habitats must have a high dispersal ability between populations as they would not be able to survive without the ability to move between their patchily distributed habitats (Arribas et al. 2012; Hof et al. 2012). Therefore, species inhabiting lentic water habitats tend to have relatively high dispersal ability, a wide distribution, and exhibit little genetic structuring between geographical regions.Given that wet rock habitats exist as isolated patches within river systems and are tend to disappearance (Wantzen and Junk 2009), it is likely that species adapted to these environments require high dispersal ability. In this study, we compared congeneric species within *Dolophilodes* and *Cryptoperla*, and revealed low levels of genetic differentiation in species adapted to wet rock habitats, supporting the possibility that this may be a general trend. Also, in this study, *Do. angustata* adapted to wet rock habitats was occasionally detected in flowing water environments, when large bedrock surfaces were present in upstream areas. However, the reverse was not observed. Although it remains unclear to what extent these species utilize bedrock within the main stream, such environments are rarely continuous or widespread. Therefore, even when these habitats are accessible, they can still be considered spatially discrete, point-like environments.

These habitats have received little focus in river ecology (Wantzen and Junk 2009; Girón et al. 2021). However, in these wet rock habitats we observed the target species of this study, *Dolophilodes* and *Cryptoperla*and *Baetis thermicus* (Ephemeroptera), *Nemoura* (Plecoptera), Rhyacophilidae (Trichoptera), and Dytiscidae (Coleoptera). This suggests that wet rock habitats represent an important component of river ecosystems (Wantzen and Junk 2009; Girón et al. 2021).

For the genus *Dolophilodes*, which includes species inhabiting wet rock habitats, headwater species *Do.japonica* and upstream *species Do. commata* showed that species inhabiting headwater habitats exhibited greater genetic differentiation and more complex genetic structures. A similar pattern was also observed in *C. kawasawai*. In contrast, *C. japonica*, a congeneric species that inhabits upstream areas, showed almost no genetic differentiation among sites, compared with upstream species of other genera. Factors other than environmental conditions may underlie this pattern; however, we were unable to identify them in the present study. Compared with *C. kawasawai*, *C. japonica* showed a slightly higher level of genetic differentiation. Overall, our results indicate that the spatial arrangement of habitats to which species are adapted strongly shapes population genetic structure. In combination with species’ dispersal ability, it also determines the applicable models of dispersal capacity (Hughes 2009; 2013; Phillipsen et al. 2015). By comparing congeneric species, this study not only supports previous conclusions but also clarifies that wet rock habitats, although part of riverine ecosystems, exhibit a spatial configuration and genetic pattern that are analogous to those of lentic habitats.

In conclusion, there are very few studies that have compared the genetic structure across different environments among different species of the same genus. When comparisons are limited to a single taxonomic group, it becomes difficult to rule out patterns that may be specific to that group. In this study, we demonstrated this pattern using multiple genera in which closely related species replace each other along the river course continuum. This pattern is not limited to upstream vs. downstream or lotic vs. lentic environmental factors. Genetic structure has been shown to vary with elevation (Giordano et al. 2007; Mikami et al. 2023; Suzuki et al. 2024) and to reflect differences in gene flow (Wolcock et al. 2007; Suzuki et al. 2024). Species adapted to high-elevation zones tend to exhibit genetic differentiation among mountain regions, as lower elevation areas between act as barriers to dispersal (Uscanga et al. 2021; Mikami et al. 2023; Suzuki et al. 2024).

It can be inferred that species adapted to wet rock habitats have survived to the present due to their high dispersal ability. However, the ecology of wet rock habitats remains largely unexplored, and further studies involving a broader range of species are needed. This finding is novel and represents a new insight into habitat-driven genetic structure in river ecosystems.

## Acknowledgements

We express our thanks to the Muroto Global Geopark Center for supporting our investigation. This study conducted English proofreading using M365 Copilot, which utilizes OpenAI’s language models. We express our thanks to Shimura, N., Kuhara, N. for advice about species information. This study was supported by Muroto UNESCO Global Geopark Research Grant 2018, 2020 (M.T.).

## Supporting Information

Figure S1 The environments surveyed in this study are arranged for each river from left to right as wet rock, and from headwater to downstream.

Figure S2 Box-plot showing range of environmental factors in each cluster. The range of altitude in each group (B); the range of riverbed slope degree in each group (B); the range of channel width in each group (C); the range of canopy openness in each group (D); the range of substrate coarseness in each group (E)

Figure S3 Haplotype networks based on the mtDNA COI region (## bp) for *Dipteromimus tipuliromis*, *Isonychia japonica*, and *Stenopsyche marmorata*. The colors of circles correspond to the results of hierarchal clustering in figure 3 as follows: group 1 is headwater environments and group 2 is upstream environments, group 3 is middle stream environments, group 4 is downstream environments.

## Data Availability Statement

The DNA sequences have been deposited in the public repository of GenBank systems (accession numbers: LC904292- LC904613). All methods information is included in this manuscript. All specimens used in this study are stored in the Tojo laboratory of Shinshu University.

## References

Alp M, Keller I, Westram AM, Robinson CT (2012) How river structure and biological traits influence gene flow: a population genetic study of two stream invertebrates with differing dispersal abilities. Freshw Biol 57:969– 981. 10.1111/j.1365-2427.2012.02758.x

de Araujo Barbosa V, Graham SE, Hogg ID, Smith BJ, McGaughran A (2025) A landscape genetics approach reveals species-specific connectivity patterns for stream insects in fragmented habitats. Ecol Evol 15:e71084. 10.1002/ece3.71084

Arribas P, Velasco J, Abellán P, Sánchez-Fernández D, et al (2012) Dispersal ability rather than ecological tolerance drives differences in range size between lentic and lotic water beetles (Coleoptera: Hydrophilidae). J Biogeogr 39:984–994. 10.1111/j.1365-2699.2011.02641.x

Bain MB, Finn JT, Booke HE (1985) Quantifying stream substrate for habitat analysis studies. N Am J Fish Manag 5:499–506. 10.1577/1548-8659(1985)5<499:QSSFHA>2.0.CO;2

Baggiano O, Schmidt DJ, Sheldon F, Hughes JM (2011) The role of altitude and associated habitatstability in determining patterns of population genetic structure in two species of *Atalophlebia* (Ephemeroptera: Leptophlebiidae). Freshw Biol 56:230–249. 10.1111/j.1365-2427.2010.02490.x

Bowler DA, Benton TG (2005) Causes and consequences of animal dispersal strategies: relating individual behaviour to spatial dynamics. Biol Rev 8:205–225. 10.1017/S1464793104006645

Campbell RE, McIntosh AR (2013) Do isolation and local habitat jointly limit the structure of stream invertebrate assemblages? Freshw Biol 58:128–141. 10.1111/fwb.12045

Clement M, Posada DC, Crandall KA (2000) TCS: a computer program to estimate gene genealogies. Mol Ecol 9:1657–1659. 10.1046/j.1365-294x.2000.01020.x

Davis CD, Epps CW, Flitcroft RL, Banks MA (2018) Refining and defining riverscape genetics: How rivers influence population genetic structure. Wiley Interdiscip Rev Water 5:e1269. 10.1002/wat2.1269

Doretto A, Piano E, Larson CE (2020) The river continuum cncept: lessons from the past and perspectives for the future. Can J Fish Aquat Sci 77:1853–1864. 10.1139/cjfas-2020-0039

Finn DS, Adler PH (2006) Population genetic structure of a rare high-elevation black fly, *Metacnephia coloradensis*, occupying Colorado lake outlet streams. Freshw Biol 51:2240–2251. 10.1111/j.1365-2427.2006.01647.x

Finn DS, Blouin MS, Lytle DA (2007) Population genetic structure reveals terrestrial affinities for a headwater stream insect. Freshw Biol 52:1881–1897. 10.1111/j.1365-2427.2007.01813.x

Folmer O, Black M, Hoeh W, Lutz R, Vrijenhoek R (1994) DNA primers for amplification of mitochondrial cytochrome c oxidase subunit I from diverse metazoan invertebrates. Mol Mar Biol Biotechnolog 3:294–299.

Fusco NA, Pehek E, Munshi-South J (2021) Urbanization reduces gene flow but not genetic diversity of stream salamander populations in the New York City metropolitan area. Evol Appl 14:99–116. 10.1111/eva.13025

Giordano AR, Ridenhour BJ, Storfer A (2007) The influence of altitude and topography on genetic structure in the long-toed salamander (Ambystoma macrodactulym). Mol Ecol 16:1625–1637. 10.1111/j.1365-294X.2006.03223.x

Girón JC, Short AEZ (2021) The Acidocerinae (Coleoptera, Hydrophilidae): taxonomy, classification, and catalog of species. ZooKeys 1045:1–236. 10.3897/zookeys.1045.63810

Hjalmarsson AE, Bergsten J, Monaghan MT (2015) Dispersal is linked to habitat use in 59 species of water beetles (Coleoptera: Adephaga) on Madagascar. Ecography 38:732–739. 10.1111/ecog.01138

Hof C, Brändle M, Dehling DM, et al (2012) Habitat stability affects dispersal and the ability to track climate change. Biol Lett 8:639–643. 10.1098/rsbl.2012.0023

Hughes JM, Schmidt DJ, Finn DS (2009) Genes in streams: using DNA to understand the movement of freshwater fauna and their riverine habitat. BioScience 59:573–583. 10.1525/bio.2009.59.7.8

Hughes JM, Huey JA, Schmidt DJ (2013) Is realised connectivity among populations of aquatic fauna predictable from potential connectivity? Freshw Biol 58:951–966. 10.1111/fwb.12099

Ikeda H, Nishikawa M, Sota T (2012) Loss of flight promotes beetle diversification. Nat Commun 3:1–8. 10.1038/ncomms1659

Katoh K, Standley DM (2013) MAFFT multiple sequence alignment software version 7: improvements in performance and usability. Mol Biol Evol 30:772–780. 10.1093/molbev/mst010

Kindlmann P, Burel F (2008) Connectivity measures: a review. Landsc Ecol 23:879–890. 10.1007/s10980-008-9245-4

Kawai T, Tanida K (2018) Aquatic Insects of Japan: Manual with Keys and Illustrations. The second edition. Tokai University, Kanagawa

Kuhara N (2005) Taxonomic revision of the genus *Dolophilodes* subgenus *Dolophilodes* (Trichoptera: Philopotamidae) of Japan. Entomol Sci 8:91–107. 10.1111/j.1479-8298.2005.00104.x

Kuhara N (2018) Philopotamidae. In: Kawai T, Tanida K (eds) Aquatic insects of Japan: manual with keys and illustrations, 2nd edn. Tokai University Press, pp 529–543

Kumar S, Stecher G, Tamura K (2016) MEGA7: molecular evolutionary genetics analysis version 7.0 for bigger datasets. Mol Biol Evol 33:1870–1874. 10.1007/s10980-008-9245-4

Lancaster J, Downes BJ, Finn DS, St Clair RM (2024) Connected headwaters: indelible field evidence of dispersal by a diverse caddisfly assemblage up stream valleys to dry catchment boundaries. Freshw Biol 69:1–14. 10.1111/fwb.14188

Leigh JW, Bryant D, Nakagawa S (2015) POPART: full-feature software for haplotype network construction. Methods Ecol Evol 6:1110–1116. 10.1111/2041-210X.12410

Mikami K, Takenaka M, Nozaki T, Bae YJ, Tojo K (2023) Phylogeography of alpine and subalpine adapted *Pseudostenophylax* caddisflies (Limnephilidae: Trichoptera): a strong relationship with mountain formation. Biol J Linn Soc 139:257–274. 10.1093/biolinnean/blad022

Miller 2005

Miyazono S, Taylor CM (2013) Effects of habitat size and isolation on species immigration–extinction dynamics and community nestedness in a desert river system. Freshw Biol 58:1303–1312. 10.1111/fwb.12127

Monaghan MT, Spaak P, Robinson CT, Ward JV (2002) Population genetic structure of 3 alpine stream insects: influences of gene flow, demographics, and habitat fragmentation. J-NABS 21:114–131.

Murray BF, Reid MA, Capon SJ, Thoms M, Wu SB (2019) Gene flow and genetic structure in *Acacia stenophylla* (Fabaceae): Effects of hydrological connectivity. J Biogeogr 46:1138–1151. 10.1111/jbi.13566

Ogitani M, Nakamura H (2008) Distribution and seasonal population change of Heptageniidae nymph in the Oguro River (The branch of Tenryu River). Ann Environ Sci Shinshu Univ 30:57–66.

Ohba SY, Suzuki T, Fukui M, et al (2025) Flight characteristics and phylogeography in three large-bodied diving beetle species: evidence that the species with expanded distribution is an active flier. Biol J Linn Soc blae017. 10.1093/biolinnean/blae017

Ohnishi O, Takenaka M, Okano R, Yoshitomi H, Tojo K (2021) Wide-scale gene flow, even in insects that have lost their flight ability: presence of dispersion due to a unique parasitic ecological strategy of piggybacking hosts. Zool Sci 38:122–139. 10.2108/zs200088

Okamoto S, Tojo K (2021) Distribution patterns and niche segregation of three closely related Japanese ephemerid mayflies: a re-examination of each species’ habitat from “megadata” held in the “National Census on River Environments”. Limnology 22:277–287. 10.1007/s10201-021-00654-2

Okamoto S, Saito T, Tojo K (2022) Geographical fine-scaled distributional differentiation caused by niche differentiation in three closely related mayflies. Limnology 23:89–101. 10.1007/s10201-021-00673-z

Pauls SU, Lumbsch HT, Haase P (2006) Phylogeography of the montane caddisfly *Drusus discolor*: evidence for multiple refugia and periglacial survival. Mol Ecol 15:2153–2169. 10.1111/j.1365-294X.2006.02916.x

Paz-Vinas I, Loot G, Stevens VM, Blanchet S (2015) Evolutionary processes driving spatial patterns of intraspecific genetic diversity in river ecosystems. Mol Ecol 24:4586–4604. 10.1111/mec.13345

Phillipsen IC, Kirk EH, Bogan MT, et al (2015) Dispersal ability and habitat requirements determine landscape-level genetic patterns in desert aquatic insects. Mol Ecol 24:54–69. 10.1111/mec.13003

Prendini L, Weygoldt P, Wheeler WC (2005) Systematics of the Damon variegatus group of African whip spiders (Chelicerata: Amblypygi): evidence from behaviour, morphology and DNA. Org Divers Evol 5:203–236. 10.1016/j.ode.2004.12.004

R Core Team (2020) R: a language and environment for statistical computing. R Foundation for Statistical Computing, Vienna

Rozas J, Sánchez-DelBarrio JC, Messeguer X, Rozas R (2003) DnaSP, DNA polymorphism analyses by the coalescent and other methods. Bioinformatics 19:2496–2497. 10.1093/bioinformatics/btg359

Saito R, Tojo K (2016) Comparing spatial patterns of population density, biomass, and genetic diversity patterns of the habitat generalist mayfly *Isonychia japonica* Ulmer (Ephemeroptera: Isonychiidae) in the Chikuma– Shinano River basin. Freshw Sci 35:724–737. 10.1086/686537

Sexton JP, Hangartner SB, Hoffmann AA (2014) Genetic isolation by environment or distance: which pattern of gene flow is most common? Evolution 68:1–15. 10.1111/evo.12258

Sproul JS, Houston DD, Davis N, et al (2014) Comparative phylogeography of codistributed aquatic insects in western North America: insights into dispersal and regional patterns of genetic structure. Freshw Biol 59:2051–2063. 10.1111/fwb.12406

Suzuki H, Takenaka M, Tojo K (2024) Evolutionary history of a cold-adapted limnephilid caddisfly: Effects of climate change and topography on genetic structure. Mol Phylogenet Evol 191:107967. 10.1016/j.ympev.2023.107967

Takenaka M, Tojo K (2019) Ancient origin of a dipteromimid mayfly family endemic to the Japanese Islands and its genetic differentiation across tectonic faults. Biol J Linn Soc 126:555–573. 10.1093/biolinnean/bly192

Takenaka M, Tokiwa T, Tojo K (2019) Concordance between molecular biogeography of *Dipteromimus tipuliformis* and geological history in the local fine scale (Ephemeroptera, Dipteromimidae). Mol Phylogenet Evol 139:106547. 10.1016/j.ympev.2019.106547

Takenaka M, Shibata S, Ito T, Shimura N, Tojo K (2021) Phylogeography of the northernmost distributed *Anisocentropus* caddisflies and their comparative genetic structures based on habitat preferences. Ecol Evol 11:4957–4971. 10.1002/ece3.7419

Tyers M (2017) riverdist: River Network Distance Computation and Applications. R package version 0.15.0. https://CRAN.R-project.org/package=riverdist

Ueki G, Tojo K (2023) The phylogeography of the stag beetle *Dorcus montivagus* (Coleoptera, Lucanidae): Comparison with the phylogeography of its specific host tree, the Japanese beech Fagus crenata. Entomol Sci 26:e12535. 10.1111/ens.12535

Uscanga A, López H, Piñero D, Emerson BC, Mastretta-Yanes A (2021) Evaluating species origins within tropical sky-islands arthropod communities. J Biogeogr 48:2199–2210. 10.1111/jbi.14144

Vannote RL, Minshall GW, Cummins KW, Sedell JR, Cushing CE (1980) The River Continuum Concept. Can J Fish Aquat Sci 37:130–137. 10.1139/f80-017

Wantzen KM, Junk WJ (2009) Riparian Wetlands. In: Jørgensen SE (ed) Ecosystem ecology. Academic Press, pp 342–351

Waters JM, Craw D, Burridge CP, et al (2015) Within-river genetic connectivity patterns reflect contrasting geomorphology. J Biogeogr 42:2452–2460. 10.1111/jbi.12608

Waters JM, Emerson BC, Arribas P, McCulloch GA (2020) Dispersal reduction: Causes, genomic mechanisms, and evolutionary consequences. Trends Ecol Evol 35:512–522. 10.1016/j.tree.2020.01.012

Waters JM, King TM, Craw D (2024) Gorges partition diversity within New Zealand flathead Galaxias populations. J Fish Biol 104:950–956. 10.1111/jfb.15635

Wright S (1943) Isolation by distance. Genetics 28:114–138. 10.1093/genetics/28.2.114

Yoshikawa N, Matsui M, Nishikawa K, Kim JB, Kryukov A (2008) Phylogenetic relationships and biogeography of the Japanese clawed salamander, *Onychodactylus japonicus* (Amphibia: Caudata: Hynobiidae), and its congener inferred from the mitochondrial cytochrome b gene. Mol Phylogenet Evol 49:249–259. 10.1016/j.ympev.2008.07.016

Zhou J, Kang S, Schadt CW, Garten CT Jr (2008) Spatial scaling of functional gene diversity across various microbial taxa. Proc Natl Acad Sci 105:7768–7773. 10.1073/pnas.0709016105

