## Supplemental files for "Comparing the Influence of Habitat Configuration on Population Connectivity and Genetic Structure Using Congeneric Species Across Multiple Taxa"

| Table S1 Sampling sites | |  |  |  |
| --- | --- | --- | --- | --- |
|  | Sites | Rivers | GPS |  |
| st. 5 | Sakihamacho, Muroto, Kochi | Higashino River | 33.39079 | 134.16807 |
| st. 6 | Sakihamacho, Muroto, Kochi | Higashino River | 33.38827 | 134.16728 |
| st. 7 | Kiragawa, Muroto, Kochi | Higashino River | 33.33238 | 134.10399 |
| st. 8 | Kiragawa, Muroto, Kochi | Higashino River | 33.34629 | 134.11847 |
| st. 9 | Kiragawa, Muroto, Kochi | Higashino River | 33.36003 | 134.13502 |
| st. 10 | Kiragawa, Muroto, Kochi | Higashino River | 33.37386 | 134.14661 |
| st. 11 | Kiragawa, Muroto, Kochi | Higashino River | 33.38250 | 134.15898 |
| st. 12 | Kiragawa, Muroto, Kochi | Higashino River | 33.38250 | 134.15898 |
| st. 13 | Kiragawa, Muroto, Kochi | Higashino River | 33.35037 | 134.12195 |
| st. 14 | Hirose, Kitagawa, Kochi | Nahari River | 33.49733 | 134.15916 |
| st. 15 | Hirose, Kitagawa, Kochi | Nahari River | 33.49809 | 134.15813 |
| st. 16 | Hirose, Kitagawa, Kochi | Nahari River | 33.50775 | 134.14811 |
| st. 17 | Notomoko, Kitagawa, Kochi | Nahari River | 33.44159 | 134.03174 |
| st. 18 | Tochinoki, Aki, Kochi | Aki River | 33.55626 | 133.90126 |
| st. 19 | Minato, Aki, Kochi | Aki River | 33.50014 | 133.91368 |
| st. 20 | Akinokawako, Aki, Kochi | Aki River | 33.57844 | 133.92364 |
| st. 21 | Monobe, Kami, Kochi | Aki River | 33.64996 | 133.90922 |
| st. 22 | Monobe, Kami, Kochi | Aki River | 33.64996 | 133.90922 |
| st. 23 | Maikawa, Aki, Kochi | Aki River | 33.64146 | 133.90360 |
| st. 24 | Maikawa, Aki, Kochi | Aki River | 33.64030 | 133.90267 |
| st. 25 | Maikawa, Aki, Kochi | Aki River | 33.63943 | 133.90431 |
| st. 26 | Akinokawaotsu, Aki, Kochi | Aki River | 33.59821 | 133.91046 |
| st. 27 | Kagami, Konan, Kochi | Monobe River | 33.64045 | 133.84511 |
| st. 28 | Kagami, Konan, Kochi | Monobe River | 33.64045 | 133.84511 |
| st. 29 | Monobe, Kami, Kochi | Monobe River | 33.66750 | 133.88338 |
| st. 30 | Monobe, Kami, Kochi | Monobe River | 33.68618 | 133.86339 |
| st. 31 | Kahoku, Kami, Kochi | Monobe River | 33.68561 | 133.84378 |
| st. 32 | Noichi, Konan, Kochi | Monobe River | 33.56604 | 133.68153 |
| st. 33 | Monobe, Kami, Kochi | Monobe River | 33.55289 | 133.68080 |
| st. 34 | Oi, Kaiyo, Tokushima | Kaifu River | 33.62897 | 134.31721 |
| st. 35 | Ogawa, Kaiyo, Tokushima | Kaifu River | 33.70552 | 134.30269 |
| st. 36 | Ogawa, Kaiyo, Tokushima | Kaifu River | 33.71156 | 134.30685 |
| st. 37 | Ogawa, Kaiyo, Tokushima | Kaifu River | 33.71156 | 134.30685 |
| st. 38 | Ogawa, Kaiyo, Tokushima | Kaifu River | 33.70782 | 134.30696 |
| st. 39 | Ogawa, Kaiyo, Tokushima | Kaifu River | 33.72628 | 134.29247 |
| st. 40 | Kaikawa, Naka, Tokushima | Naka River | 33.75247 | 134.26547 |
| st. 41 | Kaikawa, Naka, Tokushima | Naka River | 33.75787 | 134.26046 |
| st. 42 | Kaikawa, Naka, Tokushima | Naka River | 33.77821 | 134.25483 |
| st. 43 | Yananoe, Naka, Tokushima | Naka River | 33.82650 | 134.48901 |
| st.101 | Nakagawa, Anan, Tokushima | Monobe River | 33.939978 | 134.644304 |
| st.102 | Hiura, Naka, Tokushima | Monobe River | 33.801787 | 134.400524 |
| st.104 | Kaikawa, Naka, Tokushima | Monobe River | 33.776489 | 134.237711 |
| st.106 | Ogawa, Kaiyo, Tokushima | Kaifu River | 33.725318 | 134.301261 |
| st.108 | Kagami, Konan, Kochi | Monobe River | 33.625989 | 133.848973 |
| st.109 | Kagami, Konan, Kochi | Monobe River | 33.625989 | 133.848973 |
| st.110 | Kagami, Konan, Kochi | Monobe River | 33.625989 | 133.848973 |
| st.111 | Kagami, Konan, Kochi | Monobe River | 33.657281 | 133.871325 |
| st.113 | Monobe, Kami, Kochi | Monobe River | 33.675215 | 133.873643 |
| st.117 | Akinokawako, Aki, Kochi | Aki River | 33.583948 | 133.928695 |
| st.118 | Akinokawako, Aki, Kochi | Aki River | 33.581176 | 133.924982 |
| st.119 | Akinokawako, Aki, Kochi | Aki River | 33.574285 | 133.925016 |
| st.120 | Akinokawako, Aki, Kochi | Aki River | 33.569221 | 133.919276 |
| st.122 | Hirose, Kitagawa, Kochi | Nahari River | 33.497330 | 134.159160 |
| st.124 | Hirose, Kitagawa, Kochi | Nahari River | 33.513245 | 134.135219 |
| st.125 | Hirose, Kitagawa, Kochi | Nahari River | 33.505501 | 134.147646 |
| st.127 | Ogawa, Kaiyo, Tokushima | Kaifu River | 33.682683 | 134.305579 |
| st.128 | Ogawa, Kaiyo, Tokushima | Kaifu River | 33.702145 | 134.302768 |
| st.129 | Akinokawako, Aki, Kochi | Higashino River | 33.345245 | 134.118011 |
| st.130 | Akinokawako, Aki, Kochi | Higashino River | 33.389157 | 134.167209 |
| st.131 | Akinokawako, Aki, Kochi | Higashino River | 33.389157 | 134.167209 |
| st.132 | Akinokawako, Aki, Kochi | Higashino River | 33.377373 | 134.154056 |
| st.133 | Akinokawako, Aki, Kochi | Higashino River | 33.339317 | 134.110432 |
| st.135 | Akinokawako, Aki, Kochi | Higashino River | 33.370292 | 134.131502 |
| Higashino river (drainage basin: 21.5 km^2^; main channel length: 9.2 km; number of sampling site: 15 sites), Nahari river (drainage basin: 311.3 km^2^; main channel length: 56.1 km; number of sampling site: seven sites), Aki river (drainage basin: 143.5 km^2^; main channel length: 27.8 km, number of sampling site: 13 sites), Monobe river (drainage basin: 508.0 km^2^; main channel length: 290.0 km, number of sampling site: 13 sites), Naka river (drainage basin: 874.0 km^2^; main channel length: 125.0 km, number of sampling site: six sites), Kaifu river (drainage basin: 206.0 km^2^; main channel length: 36.0 km, number of sampling site: nine sites) | | | | |

| Table S2 List of specimens examined in this study and the GenBank accession numbers | | | | | |
| --- | --- | --- | --- | --- | --- |
|  |  |  | GPS |  |  |
| Species Name | Sampling sites | | Latitude | Longitude | GenBank acces. No. |
| *Ephemera japonica* | | |  |  |  |
|  | st. 26 | Akinokawaotsu, Aki, Kochi | 33.598210 | 133.910460 | LC904292 |
|  | st. 21 | Monobe, Kami, Kochi | 33.649960 | 133.909220 | LC904293 |
|  | st. 21 | Monobe, Kami, Kochi | 33.649960 | 133.909220 | LC904294 |
|  | st. 21 | Monobe, Kami, Kochi | 33.649960 | 133.909220 | LC904295 |
|  | st. 21 | Monobe, Kami, Kochi | 33.649960 | 133.909220 | LC904296 |
|  | st. 21 | Monobe, Kami, Kochi | 33.649960 | 133.909220 | LC904297 |
|  | st. 37 | Ogawa, Kaiyo, Tokushima | 33.711560 | 134.306850 | LC904298 |
|  | st. 37 | Ogawa, Kaiyo, Tokushima | 33.711560 | 134.306850 | LC904299 |
|  | st. 37 | Ogawa, Kaiyo, Tokushima | 33.711560 | 134.306850 | LC904300 |
|  | st. 37 | Ogawa, Kaiyo, Tokushima | 33.711560 | 134.306850 | LC904301 |
|  | st. 37 | Ogawa, Kaiyo, Tokushima | 33.711560 | 134.306850 | LC904302 |
|  | st. 41 | Kaikawa, Naka, Tokushima | 33.757870 | 134.260490 | LC904303 |
|  | st. 41 | Kaikawa, Naka, Tokushima | 33.757870 | 134.260490 | LC904304 |
|  | st. 41 | Kaikawa, Naka, Tokushima | 33.757870 | 134.260490 | LC904305 |
|  | st. 41 | Kaikawa, Naka, Tokushima | 33.757870 | 134.260490 | LC904306 |
|  | st. 41 | Kaikawa, Naka, Tokushima | 33.757870 | 134.260490 | LC904307 |
|  | st. 14 | Hirose, Kitagawa, Kochi | 33.497330 | 134.159160 | LC904308 |
|  | st. 14 | Hirose, Kitagawa, Kochi | 33.497330 | 134.159160 | LC904309 |
|  | st. 14 | Hirose, Kitagawa, Kochi | 33.497330 | 134.159160 | LC904310 |
|  | st. 14 | Hirose, Kitagawa, Kochi | 33.497330 | 134.159160 | LC904311 |
|  | st. 14 | Hirose, Kitagawa, Kochi | 33.497330 | 134.159160 | LC904312 |
|  | st. 9 | Kiragawa, Muroto, Kochi | 33.360030 | 134.135020 | LC904313 |
|  | st. 9 | Kiragawa, Muroto, Kochi | 33.360030 | 134.135020 | LC904314 |
|  | st. 9 | Kiragawa, Muroto, Kochi | 33.360030 | 134.135020 | LC904315 |
|  | st. 9 | Kiragawa, Muroto, Kochi | 33.360030 | 134.135020 | LC904316 |
|  | st. 9 | Kiragawa, Muroto, Kochi | 33.360030 | 134.135020 | LC904317 |
|  | st. 28 | Kagami, Konan, Kochi | 33.640450 | 133.845110 | LC904318 |
|  | st. 28 | Kagami, Konan, Kochi | 33.640450 | 133.845110 | LC904319 |
|  | st. 28 | Kagami, Konan, Kochi | 33.640450 | 133.845110 | LC904320 |
|  | st. 28 | Kagami, Konan, Kochi | 33.640450 | 133.845110 | LC904321 |
|  | st. 28 | Kagami, Konan, Kochi | 33.640450 | 133.845110 | LC904322 |
| *Ephemera strigata* | | |  |  |  |
|  | st. 26 | Akinokawaotsu, Aki, Kochi | 33.598210 | 133.910460 | LC904323 |
|  | st. 26 | Akinokawaotsu, Aki, Kochi | 33.598210 | 133.910460 | LC904324 |
|  | st. 26 | Akinokawaotsu, Aki, Kochi | 33.598210 | 133.910460 | LC904325 |
|  | st. 26 | Akinokawaotsu, Aki, Kochi | 33.598210 | 133.910460 | LC904326 |
|  | st. 26 | Akinokawaotsu, Aki, Kochi | 33.598210 | 133.910460 | LC904327 |
|  | st. 34 | Ogawa, Kaiyo, Tokushima | 33.628970 | 134.317210 | LC904328 |
|  | st. 34 | Ogawa, Kaiyo, Tokushima | 33.628970 | 134.317210 | LC904329 |
|  | st. 34 | Ogawa, Kaiyo, Tokushima | 33.628970 | 134.317210 | LC904330 |
|  | st. 43 | Yananoe, Naka, Tokushima | 33.826500 | 134.489010 | LC904331 |
|  | st. 43 | Yananoe, Naka, Tokushima | 33.826500 | 134.489010 | LC904332 |
|  | st. 43 | Yananoe, Naka, Tokushima | 33.826500 | 134.489010 | LC904333 |
|  | st. 43 | Yananoe, Naka, Tokushima | 33.826500 | 134.489010 | LC904334 |
|  | st. 43 | Yananoe, Naka, Tokushima | 33.826500 | 134.489010 | LC904335 |
|  | st. 17 | Notomoko, Kitagawa, Kochi | 33.441590 | 134.031740 | LC904336 |
|  | st. 17 | Notomoko, Kitagawa, Kochi | 33.441590 | 134.031740 | LC904337 |
|  | st. 17 | Notomoko, Kitagawa, Kochi | 33.441590 | 134.031740 | LC904338 |
|  | st. 17 | Notomoko, Kitagawa, Kochi | 33.441590 | 134.031740 | LC904339 |
|  | st. 17 | Notomoko, Kitagawa, Kochi | 33.441590 | 134.031740 | LC904340 |
|  | st. 33 | Monobe, Kami, Kochi | 33.552890 | 133.680800 | LC904341 |
|  | st. 33 | Monobe, Kami, Kochi | 33.552890 | 133.680800 | LC904342 |
|  | st. 33 | Monobe, Kami, Kochi | 33.552890 | 133.680800 | LC904343 |
|  | st. 33 | Monobe, Kami, Kochi | 33.552890 | 133.680800 | LC904344 |
|  | st. 9 | Kiragawa, Muroto, Kochi | 33.360030 | 134.135020 | LC904345 |
|  | st. 9 | Kiragawa, Muroto, Kochi | 33.360030 | 134.135020 | LC904346 |
|  | st. 9 | Kiragawa, Muroto, Kochi | 33.360030 | 134.135020 | LC904347 |
|  | st. 9 | Kiragawa, Muroto, Kochi | 33.360030 | 134.135020 | LC904348 |
|  | st. 9 | Kiragawa, Muroto, Kochi | 33.360030 | 134.135020 | LC904349 |
| *Epeorus curvatulus* | | |  |  |  |
|  | st. 20 | Akinokawako, Aki, Kochi | 33.578440 | 133.923640 | LC904350 |
|  | st. 20 | Akinokawako, Aki, Kochi | 33.578440 | 133.923640 | LC904351 |
|  | st. 20 | Akinokawako, Aki, Kochi | 33.578440 | 133.923640 | LC904352 |
|  | st. 20 | Akinokawako, Aki, Kochi | 33.578440 | 133.923640 | LC904353 |
|  | st. 20 | Akinokawako, Aki, Kochi | 33.578440 | 133.923640 | LC904354 |
|  | st. 35 | Ogawa, Kaiyo, Tokushima | 33.705520 | 134.302690 | LC904355 |
|  | st. 35 | Ogawa, Kaiyo, Tokushima | 33.705520 | 134.302690 | LC904356 |
|  | st. 35 | Ogawa, Kaiyo, Tokushima | 33.705520 | 134.302690 | LC904357 |
|  | st. 35 | Ogawa, Kaiyo, Tokushima | 33.705520 | 134.302690 | LC904358 |
|  | st. 42 | Kaikawa, Naka, Tokushima | 33.778210 | 134.254830 | LC904359 |
|  | st. 42 | Kaikawa, Naka, Tokushima | 33.778210 | 134.254830 | LC904360 |
|  | st. 42 | Kaikawa, Naka, Tokushima | 33.778210 | 134.254830 | LC904361 |
|  | st. 42 | Kaikawa, Naka, Tokushima | 33.778210 | 134.254830 | LC904362 |
|  | st. 42 | Kaikawa, Naka, Tokushima | 33.778210 | 134.254830 | LC904363 |
|  | st. 16 | Hirose, Kitagawa, Kochi | 33.507750 | 134.148110 | LC904364 |
|  | st. 16 | Hirose, Kitagawa, Kochi | 33.507750 | 134.148110 | LC904365 |
|  | st. 16 | Hirose, Kitagawa, Kochi | 33.507750 | 134.148110 | LC904366 |
|  | st. 16 | Hirose, Kitagawa, Kochi | 33.507750 | 134.148110 | LC904367 |
|  | st. 16 | Hirose, Kitagawa, Kochi | 33.507750 | 134.148110 | LC904368 |
|  | st. 11 | Kiragawa, Muroto, Kochi | 33.382500 | 134.158980 | LC904369 |
|  | st. 11 | Kiragawa, Muroto, Kochi | 33.382500 | 134.158980 | LC904370 |
|  | st. 11 | Kiragawa, Muroto, Kochi | 33.382500 | 134.158980 | LC904371 |
|  | st. 11 | Kiragawa, Muroto, Kochi | 33.382500 | 134.158980 | LC904372 |
|  | st. 11 | Kiragawa, Muroto, Kochi | 33.382500 | 134.158980 | LC904373 |
|  | st. 32 | Noichi, Konan, Kochi | 33.566040 | 133.681530 | LC904374 |
|  | st. 32 | Noichi, Konan, Kochi | 33.566040 | 133.681530 | LC904375 |
|  | st. 32 | Noichi, Konan, Kochi | 33.566040 | 133.681530 | LC904376 |
|  | st. 32 | Noichi, Konan, Kochi | 33.566040 | 133.681530 | LC904377 |
|  | st. 32 | Noichi, Konan, Kochi | 33.566040 | 133.681530 | LC904378 |
| *Epeorus ikanonis* | | |  |  |  |
|  | st. 21 | Monobe, Kami, Kochi | 33.649960 | 133.909220 | LC904379 |
|  | st. 21 | Monobe, Kami, Kochi | 33.649960 | 133.909220 | LC904380 |
|  | st. 21 | Monobe, Kami, Kochi | 33.649960 | 133.909220 | LC904381 |
|  | st. 38 | Ogawa, Kaiyo, Tokushima | 33.707820 | 134.306960 | LC904382 |
|  | st. 38 | Ogawa, Kaiyo, Tokushima | 33.707820 | 134.306960 | LC904383 |
|  | st. 38 | Ogawa, Kaiyo, Tokushima | 33.707820 | 134.306960 | LC904384 |
|  | st. 38 | Ogawa, Kaiyo, Tokushima | 33.707820 | 134.306960 | LC904385 |
|  | st. 41 | Kaikawa, Naka, Tokushima | 33.757870 | 134.260460 | LC904386 |
|  | st. 41 | Kaikawa, Naka, Tokushima | 33.757870 | 134.260460 | LC904387 |
|  | st. 41 | Kaikawa, Naka, Tokushima | 33.757870 | 134.260460 | LC904388 |
|  | st. 41 | Kaikawa, Naka, Tokushima | 33.757870 | 134.260460 | LC904389 |
|  | st. 41 | Kaikawa, Naka, Tokushima | 33.757870 | 134.260460 | LC904390 |
|  | st.129 | Hirose, Kitagawa, Kochi | 33.345250 | 134.118010 | LC904391 |
|  | st.129 | Hirose, Kitagawa, Kochi | 33.345250 | 134.118010 | LC904392 |
|  | st.129 | Hirose, Kitagawa, Kochi | 33.345250 | 134.118010 | LC904393 |
|  | st. 31 | Kahoku, Kami, Kochi | 33.685610 | 133.843780 | LC904394 |
|  | st. 32 | Noichi, Konan, Kochi | 33.566040 | 133.681530 | LC904395 |
|  | st. 32 | Noichi, Konan, Kochi | 33.566040 | 133.681530 | LC904396 |
|  | st. 21 | Monobe, Kami, Kochi | 33.649960 | 133.909220 | LC904397 |
| *Epeorus nipponicus* | | |  |  |  |
|  | st.122 | Hirose, Kitagawa, Kochi | 33.497330 | 134.159160 | LC904399 |
|  | st.132 | Hirose, Kitagawa, Kochi | 33.497330 | 134.159160 | LC904400 |
|  | st.117 | Akinokawako, Aki, Kochi | 33.583950 | 133.928700 | LC904401 |
|  | st.117 | Akinokawako, Aki, Kochi | 33.583950 | 133.928700 | LC904402 |
|  | st.117 | Akinokawako, Aki, Kochi | 33.583950 | 133.928700 | LC904403 |
|  | st. 21 | Monobe, Kami, Kochi | 33.649960 | 133.909220 | LC904404 |
|  | st. 21 | Monobe, Kami, Kochi | 33.649960 | 133.909220 | LC904405 |
|  | st. 21 | Monobe, Kami, Kochi | 33.649960 | 133.909220 | LC904406 |
|  | st.106 | Ogawa, Kaiyo, Tokushima | 33.725318 | 134.301261 | LC904407 |
|  | st. 42 | Kaikawa, Naka, Tokushima | 33.778214 | 134.254825 | LC904398 |
|  | st. 14 | Hirose, Kitagawa, Kochi | 33.497330 | 134.159160 | LC904408 |
|  | st. 14 | Hirose, Kitagawa, Kochi | 33.497330 | 134.159160 | LC904409 |
|  | st. 14 | Hirose, Kitagawa, Kochi | 33.497330 | 134.159160 | LC904410 |
|  | st. 14 | Hirose, Kitagawa, Kochi | 33.497330 | 134.159160 | LC904411 |
|  | st. 14 | Hirose, Kitagawa, Kochi | 33.497330 | 134.159160 | LC904412 |
|  | st. 14 | Hirose, Kitagawa, Kochi | 33.497330 | 134.159160 | LC904413 |
|  | st. 11 | Kiragawa, Muroto, Kochi | 33.382500 | 134.158980 | LC904414 |
|  | st. 11 | Kiragawa, Muroto, Kochi | 33.382500 | 134.158980 | LC904415 |
|  | st. 11 | Kiragawa, Muroto, Kochi | 33.382500 | 134.158980 | LC904416 |
|  | st.108 | Kagami, Konan, Kochi | 33.625989 | 133.848973 | LC904417 |
|  | st.108 | Kagami, Konan, Kochi | 33.625989 | 133.848973 | LC904418 |
|  | st. 28 | Kagami, Konan, Kochi | 33.640450 | 133.845110 | LC904419 |
|  | st. 28 | Kagami, Konan, Kochi | 33.640450 | 133.845110 | LC904420 |
|  | st. 6 | Sakihamacho, Muroto, Kochi | 33.388274 | 134.167276 | LC904421 |
|  | st. 6 | Sakihamacho, Muroto, Kochi | 33.388274 | 134.167276 | LC904422 |
| *Dolophilodes angustata* | | |  |  |  |
|  | st. 23 | Maikawa, Aki, Kochi | 33.641460 | 133.903600 | LC904423 |
|  | st. 23 | Maikawa, Aki, Kochi | 33.641460 | 133.903600 | LC904424 |
|  | st. 23 | Maikawa, Aki, Kochi | 33.641460 | 133.903600 | LC904425 |
|  | st. 23 | Maikawa, Aki, Kochi | 33.641460 | 133.903600 | LC904426 |
|  | st. 25 | Maikawa, Aki, Kochi | 33.639430 | 133.904310 | LC904427 |
|  | st. 25 | Maikawa, Aki, Kochi | 33.639430 | 133.904310 | LC904428 |
|  | st. 25 | Maikawa, Aki, Kochi | 33.639430 | 133.904310 | LC904429 |
|  | st. 25 | Maikawa, Aki, Kochi | 33.639430 | 133.904310 | LC904430 |
|  | st. 25 | Maikawa, Aki, Kochi | 33.639430 | 133.904310 | LC904431 |
|  | st. 5 | Sakihamacho, Muroto, Kochi | 33.390790 | 134.168070 | LC904432 |
|  | st. 5 | Sakihamacho, Muroto, Kochi | 33.390790 | 134.168070 | LC904433 |
|  | st. 5 | Sakihamacho, Muroto, Kochi | 33.390790 | 134.168070 | LC904434 |
|  | st. 5 | Sakihamacho, Muroto, Kochi | 33.390790 | 134.168070 | LC904435 |
|  | st. 5 | Sakihamacho, Muroto, Kochi | 33.390790 | 134.168070 | LC904436 |
|  | st. 27 | Kagami, Konan, Kochi | 33.640450 | 133.845110 | LC904437 |
|  | st. 27 | Kagami, Konan, Kochi | 33.640450 | 133.845110 | LC904438 |
|  | st. 27 | Kagami, Konan, Kochi | 33.640450 | 133.845110 | LC904439 |
|  | st. 27 | Kagami, Konan, Kochi | 33.640450 | 133.845110 | LC904440 |
|  | st. 27 | Kagami, Konan, Kochi | 33.640450 | 133.845110 | LC904441 |
|  | st. 15 | Hirose, Kitagawa, Kochi | 33.498090 | 134.158130 | LC904442 |
|  | st. 15 | Hirose, Kitagawa, Kochi | 33.498090 | 134.158130 | LC904443 |
|  | st. 15 | Hirose, Kitagawa, Kochi | 33.498090 | 134.158130 | LC904444 |
|  | st. 15 | Hirose, Kitagawa, Kochi | 33.498090 | 134.158130 | LC904445 |
|  | st. 15 | Hirose, Kitagawa, Kochi | 33.498090 | 134.158130 | LC904446 |
|  | st. 40 | Kaikawa, Naka, Tokushima | 33.752470 | 134.265470 | LC904447 |
|  | st. 40 | Kaikawa, Naka, Tokushima | 33.752470 | 134.265470 | LC904448 |
|  | st. 40 | Kaikawa, Naka, Tokushima | 33.752470 | 134.265470 | LC904449 |
|  | st. 40 | Kaikawa, Naka, Tokushima | 33.752470 | 134.265470 | LC904450 |
|  | st. 40 | Kaikawa, Naka, Tokushima | 33.752470 | 134.265470 | LC904451 |
|  | st. 36 | Ogawa, Kaiyo, Tokushima | 33.711560 | 134.306850 | LC904452 |
|  | st. 36 | Ogawa, Kaiyo, Tokushima | 33.711560 | 134.306850 | LC904453 |
|  | st. 36 | Ogawa, Kaiyo, Tokushima | 33.711560 | 134.306850 | LC904454 |
|  | st. 36 | Ogawa, Kaiyo, Tokushima | 33.711560 | 134.306850 | LC904455 |
| *Dolophilodes commata* | | |  |  |  |
|  | st. 41 | Kaikawa, Naka, Tokushima | 33.757870 | 134.260460 | LC904456 |
|  | st. 41 | Kaikawa, Naka, Tokushima | 33.757870 | 134.260460 | LC904457 |
|  | st. 41 | Kaikawa, Naka, Tokushima | 33.757870 | 134.260460 | LC904458 |
|  | st. 41 | Kaikawa, Naka, Tokushima | 33.757870 | 134.260460 | LC904459 |
|  | st. 41 | Kaikawa, Naka, Tokushima | 33.757870 | 134.260460 | LC904460 |
|  | st. 41 | Kaikawa, Naka, Tokushima | 33.757870 | 134.260460 | LC904461 |
|  | st. 14 | Hirose, Kitagawa, Kochi | 33.497330 | 134.159160 | LC904462 |
|  | st.124 | Hirose, Kitagawa, Kochi | 33.513250 | 134.135220 | LC904463 |
|  | st.132 | Akinokawako, Aki, Kochi | 33.377370 | 134.154060 | LC904464 |
|  | st.132 | Akinokawako, Aki, Kochi | 33.377370 | 134.154060 | LC904465 |
|  | st. 29 | Monobe, Kami, Kochi | 33.667500 | 133.883380 | LC904466 |
|  | st. 29 | Monobe, Kami, Kochi | 33.667500 | 133.883380 | LC904467 |
|  | st. 29 | Monobe, Kami, Kochi | 33.667500 | 133.883380 | LC904468 |
|  | st. 29 | Monobe, Kami, Kochi | 33.667500 | 133.883380 | LC904469 |
|  | st. 29 | Monobe, Kami, Kochi | 33.667500 | 133.883380 | LC904470 |
|  | st. 29 | Monobe, Kami, Kochi | 33.667500 | 133.883380 | LC904471 |
|  | st. 29 | Monobe, Kami, Kochi | 33.667500 | 133.883380 | LC904472 |
| *Dolophilodes japonica* | | |  |  |  |
|  | st. 21 | Monobe, Kami, Kochi | 33.649960 | 133.909220 | LC904473 |
|  | st. 21 | Monobe, Kami, Kochi | 33.649960 | 133.909220 | LC904474 |
|  | st. 21 | Monobe, Kami, Kochi | 33.649960 | 133.909220 | LC904475 |
|  | st. 21 | Monobe, Kami, Kochi | 33.649960 | 133.909220 | LC904476 |
|  | st. 21 | Monobe, Kami, Kochi | 33.649960 | 133.909220 | LC904477 |
|  | st. 41 | Kaikawa, Naka, Tokushima | 33.757870 | 134.260460 | LC904478 |
|  | st. 41 | Kaikawa, Naka, Tokushima | 33.757870 | 134.260460 | LC904479 |
|  | st. 14 | Hirose, Kitagawa, Kochi | 33.497330 | 134.159160 | LC904480 |
|  | st. 14 | Hirose, Kitagawa, Kochi | 33.497330 | 134.159160 | LC904481 |
|  | st. 14 | Hirose, Kitagawa, Kochi | 33.497330 | 134.159160 | LC904482 |
|  | st. 14 | Hirose, Kitagawa, Kochi | 33.497330 | 134.159160 | LC904483 |
|  | st. 14 | Hirose, Kitagawa, Kochi | 33.497330 | 134.159160 | LC904484 |
|  | st. 14 | Hirose, Kitagawa, Kochi | 33.497330 | 134.159160 | LC904485 |
|  | st.130 | Akinokawako, Aki, Kochi | 33.389160 | 134.167210 | LC904486 |
|  | st. 28 | Kagami, Konan, Kochi | 33.640450 | 133.845110 | LC904487 |
|  | st. 28 | Kagami, Konan, Kochi | 33.640450 | 133.845110 | LC904488 |
|  | st. 28 | Kagami, Konan, Kochi | 33.640450 | 133.845110 | LC904489 |
|  | st.108 | Kagami, Konan, Kochi | 33.625990 | 133.848970 | LC904490 |
|  | st.108 | Kagami, Konan, Kochi | 33.625990 | 133.848970 | LC904491 |
|  | st.108 | Kagami, Konan, Kochi | 33.625990 | 133.848970 | LC904492 |
|  | st.108 | Kagami, Konan, Kochi | 33.625990 | 133.848970 | LC904493 |
|  | st.108 | Kagami, Konan, Kochi | 33.625990 | 133.848970 | LC904494 |
|  | st.108 | Kagami, Konan, Kochi | 33.625990 | 133.848970 | LC904495 |
|  | st.108 | Kagami, Konan, Kochi | 33.625990 | 133.848970 | LC904496 |
| *Cryptoperla japonica* | | |  |  |  |
|  | st. 11 | Kiragawa, Muroto, Kochi | 33.382500 | 134.158980 | LC904497 |
|  | st. 11 | Kiragawa, Muroto, Kochi | 33.382500 | 134.158980 | LC904498 |
|  | st. 11 | Kiragawa, Muroto, Kochi | 33.382500 | 134.158980 | LC904499 |
|  | st. 11 | Kiragawa, Muroto, Kochi | 33.382500 | 134.158980 | LC904500 |
|  | st. 11 | Kiragawa, Muroto, Kochi | 33.382500 | 134.158980 | LC904501 |
|  | st. 14 | Hirose, Kitagawa, Kochi | 33.497330 | 134.159160 | LC904502 |
|  | st. 14 | Hirose, Kitagawa, Kochi | 33.497330 | 134.159160 | LC904503 |
|  | st. 14 | Hirose, Kitagawa, Kochi | 33.497330 | 134.159160 | LC904504 |
|  | st. 14 | Hirose, Kitagawa, Kochi | 33.497330 | 134.159160 | LC904505 |
|  | st. 14 | Hirose, Kitagawa, Kochi | 33.497330 | 134.159160 | LC904506 |
|  | st. 16 | Hirose, Kitagawa, Kochi | 33.507750 | 134.148110 | LC904507 |
|  | st. 21 | Monobe, Kami, Kochi | 33.649960 | 133.909220 | LC904508 |
|  | st. 21 | Monobe, Kami, Kochi | 33.649960 | 133.909220 | LC904509 |
|  | st. 21 | Monobe, Kami, Kochi | 33.649960 | 133.909220 | LC904510 |
|  | st. 21 | Monobe, Kami, Kochi | 33.649960 | 133.909220 | LC904511 |
|  | st. 21 | Monobe, Kami, Kochi | 33.649960 | 133.909220 | LC904512 |
|  | st. 29 | Monobe, Kami, Kochi | 33.667500 | 133.883380 | LC904513 |
|  | st. 29 | Monobe, Kami, Kochi | 33.667500 | 133.883380 | LC904514 |
|  | st. 29 | Monobe, Kami, Kochi | 33.667500 | 133.883380 | LC904515 |
|  | st. 29 | Monobe, Kami, Kochi | 33.667500 | 133.883380 | LC904516 |
|  | st. 29 | Monobe, Kami, Kochi | 33.667500 | 133.883380 | LC904517 |
|  | st. 41 | Kaikawa, Naka, Tokushima | 33.757870 | 134.260460 | LC904518 |
|  | st. 41 | Kaikawa, Naka, Tokushima | 33.757870 | 134.260460 | LC904519 |
|  | st. 41 | Kaikawa, Naka, Tokushima | 33.757870 | 134.260460 | LC904520 |
|  | st. 41 | Kaikawa, Naka, Tokushima | 33.757870 | 134.260460 | LC904521 |
|  | st. 35 | Ogawa, Kaiyo, Tokushima | 33.705520 | 134.302690 | LC904522 |
|  | st. 35 | Ogawa, Kaiyo, Tokushima | 33.705520 | 134.302690 | LC904523 |
|  | st. 35 | Ogawa, Kaiyo, Tokushima | 33.705520 | 134.302690 | LC904524 |
|  | st. 35 | Ogawa, Kaiyo, Tokushima | 33.705520 | 134.302690 | LC904525 |
| *Cryptoperla kawasawai* | | |  |  |  |
|  | st. 14 | Hirose, Kitagawa, Kochi | 33.497330 | 134.159160 | LC904526 |
|  | st. 15 | Hirose, Kitagawa, Kochi | 33.498090 | 134.158130 | LC904527 |
|  | st. 23 | Maikawa, Aki, Kochi | 33.641460 | 133.903600 | LC904528 |
|  | st. 23 | Maikawa, Aki, Kochi | 33.641460 | 133.903600 | LC904529 |
|  | st. 23 | Maikawa, Aki, Kochi | 33.641460 | 133.903600 | LC904530 |
|  | st. 23 | Maikawa, Aki, Kochi | 33.641460 | 133.903600 | LC904531 |
|  | st. 23 | Maikawa, Aki, Kochi | 33.641460 | 133.903600 | LC904532 |
|  | st. 27 | Kagami, Konan, Kochi | 33.640450 | 133.845110 | LC904533 |
|  | st. 27 | Kagami, Konan, Kochi | 33.640450 | 133.845110 | LC904534 |
|  | st. 27 | Kagami, Konan, Kochi | 33.640450 | 133.845110 | LC904535 |
|  | st. 36 | Ogawa, Kaiyo, Tokushima | 33.711560 | 134.306850 | LC904536 |
|  | st. 36 | Ogawa, Kaiyo, Tokushima | 33.711560 | 134.306850 | LC904537 |
|  | st. 36 | Ogawa, Kaiyo, Tokushima | 33.711560 | 134.306850 | LC904538 |
|  | st. 36 | Ogawa, Kaiyo, Tokushima | 33.711560 | 134.306850 | LC904539 |
|  | st. 37 | Ogawa, Kaiyo, Tokushima | 33.711560 | 134.306850 | LC904540 |
|  | st. 37 | Ogawa, Kaiyo, Tokushima | 33.711560 | 134.306850 | LC904541 |
|  | st. 40 | Kaikawa, Naka, Tokushima | 33.752470 | 134.265470 | LC904542 |
|  | st. 40 | Kaikawa, Naka, Tokushima | 33.752470 | 134.265470 | LC904543 |
|  | st. 40 | Kaikawa, Naka, Tokushima | 33.752470 | 134.265470 | LC904544 |
|  | st. 40 | Kaikawa, Naka, Tokushima | 33.752470 | 134.265470 | LC904545 |
|  | st. 40 | Kaikawa, Naka, Tokushima | 33.752470 | 134.265470 | LC904546 |
|  | st. 5 | Sakihamacho, Muroto, Kochi | 33.390790 | 134.168070 | LC904547 |
|  | st. 5 | Sakihamacho, Muroto, Kochi | 33.390790 | 134.168070 | LC904548 |
|  | st. 5 | Sakihamacho, Muroto, Kochi | 33.390790 | 134.168070 | LC904549 |
|  | st. 5 | Sakihamacho, Muroto, Kochi | 33.390790 | 134.168070 | LC904550 |
| *Dipteromimus tipuliformis* | | |  |  |  |
|  | st.102 | Hiura, Naka, Tokushima | 33.801787 | 134.400524 | LC904551 |
|  | st.102 | Hiura, Naka, Tokushima | 33.801787 | 134.400524 | LC904552 |
|  | st.102 | Hiura, Naka, Tokushima | 33.801787 | 134.400524 | LC904553 |
|  | st.102 | Hiura, Naka, Tokushima | 33.801787 | 134.400524 | LC904554 |
|  | st.113 | Monobe, Kami, Kochi | 33.675215 | 133.873643 | LC904555 |
|  | st.113 | Monobe, Kami, Kochi | 33.675215 | 133.873643 | LC904556 |
|  | st.113 | Monobe, Kami, Kochi | 33.675215 | 133.873643 | LC904557 |
|  | st.119 | Akinokawako, Aki, Kochi | 33.574285 | 133.925016 | LC904558 |
|  | st.119 | Akinokawako, Aki, Kochi | 33.574285 | 133.925016 | LC904559 |
|  | st.119 | Akinokawako, Aki, Kochi | 33.574285 | 133.925016 | LC904560 |
|  | st.119 | Akinokawako, Aki, Kochi | 33.574285 | 133.925016 | LC904561 |
|  | st.135 | Akinokawako, Aki, Kochi | 33.370292 | 134.131502 | LC904562 |
|  | st.125 | Hirose, Kitagawa, Kochi | 33.505501 | 134.147646 | LC904563 |
|  | st.125 | Hirose, Kitagawa, Kochi | 33.505501 | 134.147646 | LC904564 |
|  | st.125 | Hirose, Kitagawa, Kochi | 33.505501 | 134.147646 | LC904565 |
|  | st.125 | Hirose, Kitagawa, Kochi | 33.505501 | 134.147646 | LC904566 |
|  | st.125 | Hirose, Kitagawa, Kochi | 33.505501 | 134.147646 | LC904567 |
|  | st.128 | Ogawa, Kaiyo, Tokushima | 33.702145 | 134.302768 | LC904568 |
|  | st.128 | Ogawa, Kaiyo, Tokushima | 33.702145 | 134.302768 | LC904569 |
|  | st.128 | Ogawa, Kaiyo, Tokushima | 33.702145 | 134.302768 | LC904570 |
|  | st.128 | Ogawa, Kaiyo, Tokushima | 33.702145 | 134.302768 | LC904571 |
|  | st.128 | Ogawa, Kaiyo, Tokushima | 33.702145 | 134.302768 | LC904572 |
| *Isonychia japonica* | | |  |  |  |
|  | st. 18 | Tochinoki, Aki, Kochi | 33.526260 | 133.901260 | LC904573 |
|  | st. 18 | Tochinoki, Aki, Kochi | 33.526260 | 133.901260 | LC904574 |
|  | st. 18 | Tochinoki, Aki, Kochi | 33.526260 | 133.901260 | LC904575 |
|  | st. 43 | Yananoe, Naka, Tokushima | 33.826500 | 134.489010 | LC904576 |
|  | st. 43 | Yananoe, Naka, Tokushima | 33.826500 | 134.489010 | LC904577 |
|  | st. 43 | Yananoe, Naka, Tokushima | 33.826500 | 134.489010 | LC904578 |
|  | st. 43 | Yananoe, Naka, Tokushima | 33.826500 | 134.489010 | LC904579 |
|  | st. 43 | Yananoe, Naka, Tokushima | 33.826500 | 134.489010 | LC904580 |
|  | st. 17 | Notomoko, Kitagawa, Kochi | 33.441590 | 134.031740 | LC904581 |
|  | st. 17 | Notomoko, Kitagawa, Kochi | 33.441590 | 134.031740 | LC904582 |
|  | st. 17 | Notomoko, Kitagawa, Kochi | 33.441590 | 134.031740 | LC904583 |
|  | st. 17 | Notomoko, Kitagawa, Kochi | 33.441590 | 134.031740 | LC904584 |
|  | st. 17 | Notomoko, Kitagawa, Kochi | 33.441590 | 134.031740 | LC904585 |
|  | st. 10 | Kiragawa, Muroto, Kochi | 33.373860 | 134.146610 | LC904586 |
|  | st. 8 | Kiragawa, Muroto, Kochi | 33.346290 | 134.118470 | LC904587 |
|  | st. 8 | Kiragawa, Muroto, Kochi | 33.346290 | 134.118470 | LC904588 |
|  | st. 9 | Kiragawa, Muroto, Kochi | 33.360030 | 134.135020 | LC904589 |
|  | st. 31 | Kahoku, Kami, Kochi | 33.685610 | 133.843780 | LC904590 |
| *Stenopsyche marmorata* | | |  |  |  |
|  | st. 19 | Minato, Aki, Kochi | 33.500140 | 133.913680 | LC904591 |
|  | st. 19 | Minato, Aki, Kochi | 33.500140 | 133.913680 | LC904592 |
|  | st. 19 | Minato, Aki, Kochi | 33.500140 | 133.913680 | LC904593 |
|  | st. 20 | Akinokawako, Aki, Kochi | 33.578440 | 133.923640 | LC904594 |
|  | st. 8 | Kiragawa, Muroto, Kochi | 33.346290 | 134.118470 | LC904595 |
|  | st. 8 | Kiragawa, Muroto, Kochi | 33.346290 | 134.118470 | LC904596 |
|  | st. 8 | Kiragawa, Muroto, Kochi | 33.346290 | 134.118470 | LC904597 |
|  | st. 8 | Kiragawa, Muroto, Kochi | 33.346290 | 134.118470 | LC904598 |
|  | st. 9 | Kiragawa, Muroto, Kochi | 33.360030 | 134.135020 | LC904599 |
|  | st. 9 | Kiragawa, Muroto, Kochi | 33.360030 | 134.135020 | LC904600 |
|  | st. 31 | Kahoku, Kami, Kochi | 33.685610 | 133.843780 | LC904601 |
|  | st. 17 | Notomoko, Kitagawa, Kochi | 33.441590 | 134.031740 | LC904602 |
|  | st. 17 | Notomoko, Kitagawa, Kochi | 33.441590 | 134.031740 | LC904603 |
|  | st. 16 | Hirose, Kitagawa, Kochi | 33.507750 | 134.148110 | LC904604 |
|  | st. 16 | Hirose, Kitagawa, Kochi | 33.507750 | 134.148110 | LC904605 |
|  | st. 16 | Hirose, Kitagawa, Kochi | 33.507750 | 134.148110 | LC904606 |
|  | st. 16 | Hirose, Kitagawa, Kochi | 33.507750 | 134.148110 | LC904607 |
|  | st. 43 | Yananoe, Naka, Tokushima | 33.826500 | 134.489010 | LC904608 |
|  | st. 43 | Yananoe, Naka, Tokushima | 33.826500 | 134.489010 | LC904609 |
|  | st. 43 | Yananoe, Naka, Tokushima | 33.826500 | 134.489010 | LC904610 |
|  | st. 34 | Ogawa, Kaiyo, Tokushima | 33.628970 | 134.317210 | LC904611 |
|  | st. 34 | Ogawa, Kaiyo, Tokushima | 33.628970 | 134.317210 | LC904612 |
|  | st. 34 | Ogawa, Kaiyo, Tokushima | 33.628970 | 134.317210 | LC904613 |

| Table S3 Environmental factors and sampling information in each site | | | | | | | | | | | | | |
| --- | --- | --- | --- | --- | --- | --- | --- | --- | --- | --- | --- | --- | --- |
| Site no. | | Clustering group  from Figure 3 | Altitude (m) | River bed slope degree (%) | Channel width (m) | Canopy openness (%) | Substrate coarseness | Water temperature (℃) | | EC (mS/cm) |  | DO (mg/L) |  |
|  |  |  |  |  |  |  |  | Riffle | Pool | Riffle | Pool | Riffle | Pool |
| Higashino River | |  |  |  |  |  |  |  |  |  |  |  |  |
|  | st. 5 | Wet rock | 266 | NA | NA | NA | NA | NA | NA | NA | NA | NA | NA |
|  | st. 6 | Group2 | 248 | 5.5 | 15 | 18.1 | 4.2 | 7.9 | 7 | 60 | 60.5 | 12.68 | 12.62 |
|  | st. 7 | Group3 | 4 | 1.2 | 11 | 79.7 | 3.1 | 14.6 | 14 | 81.7 | 82.6 | 11.02 | 10.89 |
|  | st. 8 | Group3 | 42 | 1.6 | 14 | 72.3 | 3.3 | 11.1 | 11.1 | 83.8 | 84.1 | 12.06 | 11.9 |
|  | st. 9 | Group2 | 87 | 0.8 | 15 | 75.3 | 4.2 | 7.2 | 6.7 | 73.2 | 72.6 | 13.97 | 13.9 |
|  | st. 10 | Group2 | 121 | 1.8 | 14 | 53.9 | 4.2 | 7.6 | 7.1 | 70 | 69.5 | 13.62 | 13.58 |
|  | st. 11 | Group2 | **177** | 3.0 | 12 | 14.4 | 4.4 | 7.6 | 7.6 | 67.4 | 67.6 | 13.11 | 13.09 |
|  | st. 12 | Group1 | **177** | 17.9 | 4 | 9.5 | 4.4 | 6 | 6 | 109.2 | 108.3 | 12.46 | 12.72 |
|  | st. 13 | Wet rock | 58 | NA | NA | NA | NA | NA | NA | NA | NA | NA | NA |
| Nahari River | |  |  |  |  |  |  |  |  |  |  |  |  |
|  | st. 14 | Group2 | 235 | 2.5 | 20 | 12.9 | 4.7 | 7 | 7 | 48.6 | 48 | 13.15 | 13.14 |
|  | st. 15 | Wet rock | 231 | NA | NA | NA | NA | NA | NA | NA | NA | NA | NA |
|  | st. 16 | Group3 | 179 | 1.5 | 65 | 73.4 | 3.8 | 6.7 | 6.7 | 57 | 56.5 | 13.31 | 13.31 |
|  | st. 17 | Group4 | 9 | 0.5 | 190 | 83.2 | 3.7 | 6.8 | 9.2 | 70.8 | 63.2 | 13.47 | 12.27 |
| Aki River | |  |  |  |  |  |  |  |  |  |  |  |  |
|  | st. 18 | Group3 | 34 | 0.6 | 88 | 97.7 | 3.8 | 7.4 | 8.3 | 102.3 | 100.9 | 13.83 | 13.78 |
|  | st. 19 | Group4 | 1 | 0.7 | 185 | 99.1 | 3.4 | 15.4 | 14.9 | 107 | 112.3 | 13.28 | 13 |
|  | st. 20 | Group3 | 71 | 0.9 | 23 | 54.6 | 4.0 | 7.3 | 7.4 | 105.3 | 105.4 | 14.27 | 14.56 |
|  | st. 21 | Group3 | **371** | 2.7 | 12.4 | 13.9 | 3.9 | 7.4 | 7.2 | 102.9 | 103 | 13.1 | 13.08 |
|  | st. 22 | Group2 | **371** | 8.5 | 4 | 12.5 | 4.5 | 8.9 | 8.1 | 87.4 | 91.8 | 12.3 | 12.3 |
|  | st. 23 | Wet rock | 313 | NA | NA | NA | NA | NA | NA | NA | NA | NA | NA |
|  | st. 24 | Group2 | 304 | 3.8 | 14 | 26.4 | 4.3 | 6.5 | 6.4 | 107.3 | 107.6 | 13.26 | 13.24 |
|  | st. 25 | Wet rock | 299 | NA | NA | NA | NA | NA | NA | NA | NA | NA | NA |
|  | st. 26 | Group3 | 122 | 1.2 | 25 | 72.4 | 3.7 | 7 | 7 | 108.8 | 111.4 | 13.41 | 13.3 |
| Monobe River | |  |  |  |  |  |  |  |  |  |  |  |  |
|  | st. 27 | Wet rock | 454 | NA | NA | NA | NA | NA | NA | NA | NA | NA | NA |
|  | st. 28 | Group2 | 454 | 6.1 | 14.2 | 10.4 | 4.0 | 6 | 5.5 | 93.4 | 92.2 | 14.14 | 14.09 |
|  | st. 29 | Group3 | 268 | 1.1 | 25 | 25.9 | 3.7 | 9.2 | 9.6 | 137.9 | 147.6 | 13.96 | 13.45 |
|  | st. 30 | Wet rock | 206 | NA | NA | NA | NA | NA | NA | NA | NA | NA | NA |
|  | st. 31 | Group3 | 104 | 0.5 | 80 | 74.0 | 3.6 | NA | NA | NA | NA | NA | NA |
|  | st. 32 | NA | 12 | 0.4 | 260 | NA | NA | NA | NA | NA | NA | NA | NA |
|  | st. 33 | Group4 | 6 | 0.3 | 262 | 100.0 | 3.4 | NA | NA | NA | NA | NA | NA |
| Kaifu River | |  |  |  |  |  |  |  |  |  |  |  |  |
|  | st. 34 | Group3 | 17 | 0.3 | 113 | 86.7 | 3.6 | 11.4 | 13 | 63.1 | 60.6 | 12.61 | 11.5 |
|  | st. 35 | Group3 | 78 | 1.0 | 53 | 68.3 | 4.2 | 5.8 | 6 | 77.6 | 78 | 14.24 | 14.25 |
|  | st. 36 | Wet rock | 133 | NA | NA | NA | NA | NA | NA | NA | NA | NA | NA |
|  | st. 37 | Group1 | 133 | 15.2 | 3.6 | 11.7 | 4.4 | 5.8 | 5.8 | 72 | 74.3 | 13.7 | 13.59 |
|  | st. 38 | Group3 | 82 | 1.2 | 21 | 34.6 | 4.0 | 6.8 | 6.4 | 82.6 | 83.7 | 13.83 | 13.87 |
|  | st. 39 |  | 518 | - | - | - | - | - | - | - | - | - | - |
| Naka River | |  |  |  |  |  |  |  |  |  |  |  |  |
|  | st. 40 | Wet rock | 496 | NA | 4 | 14.8 | 4.6 | NA | NA | NA | NA | NA | NA |
|  | st. 41 | Group2 | 454 | 1.9 | 17 | 19.4 | 4.2 | 3.7 | 3.3 | 69.2 | 74.4 | 14.41 | 14.36 |
|  | st. 42 | Group3 | 334 | 0.8 | 23 | 49.4 | 3.7 | 4.6 | 4.5 | 89.8 | 88.7 | 14.24 | 14.14 |
|  | st. 43 | Group4 | 57 | 0.4 | 140 | 96.6 | 2.5 | 6 | 6.3 | 114.5 | 113.2 | 13.96 | 13.92 |

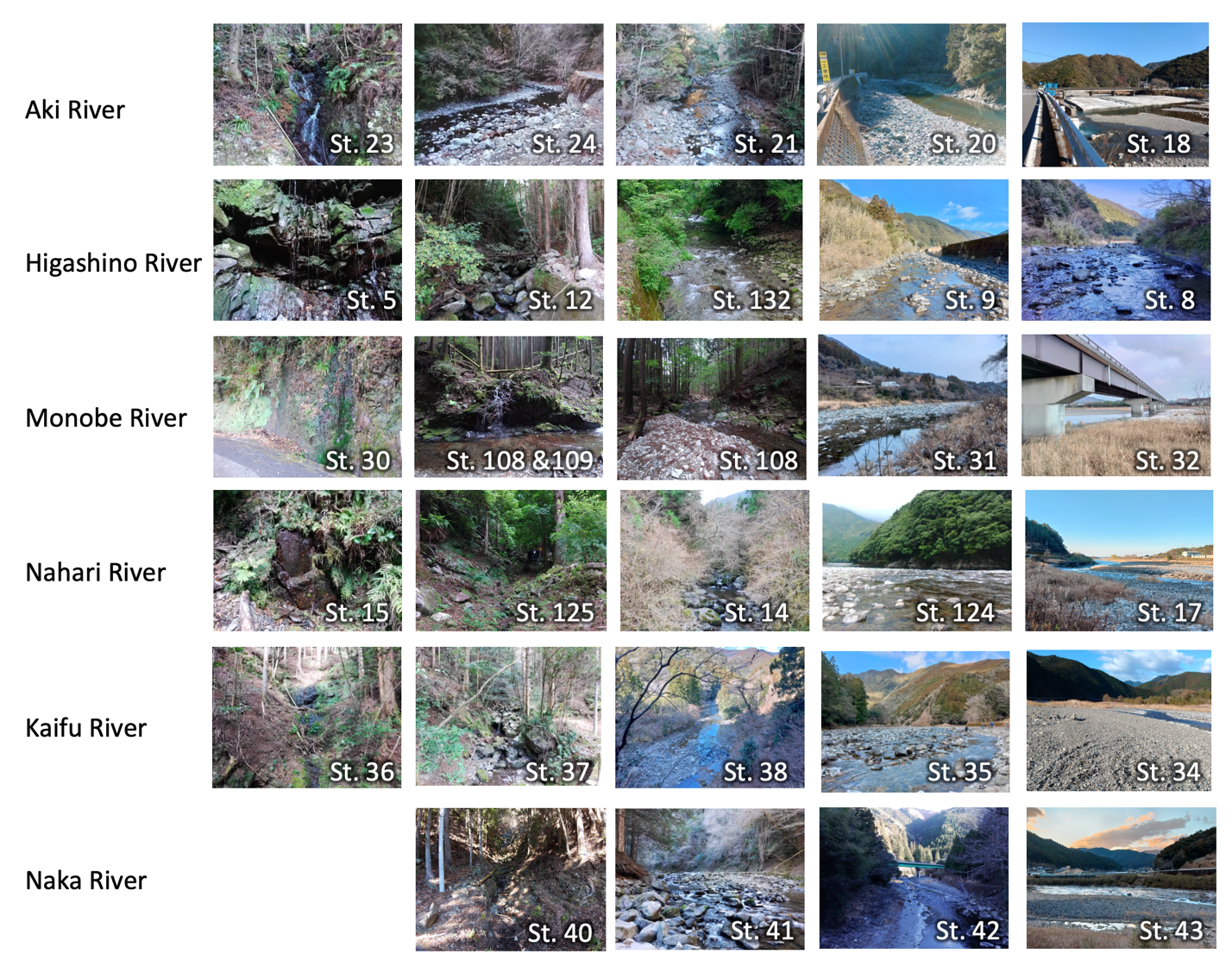

Figure S1. The environments surveyed in this study are arranged for each river from left to right as wet rock, and from headwater to downstream.

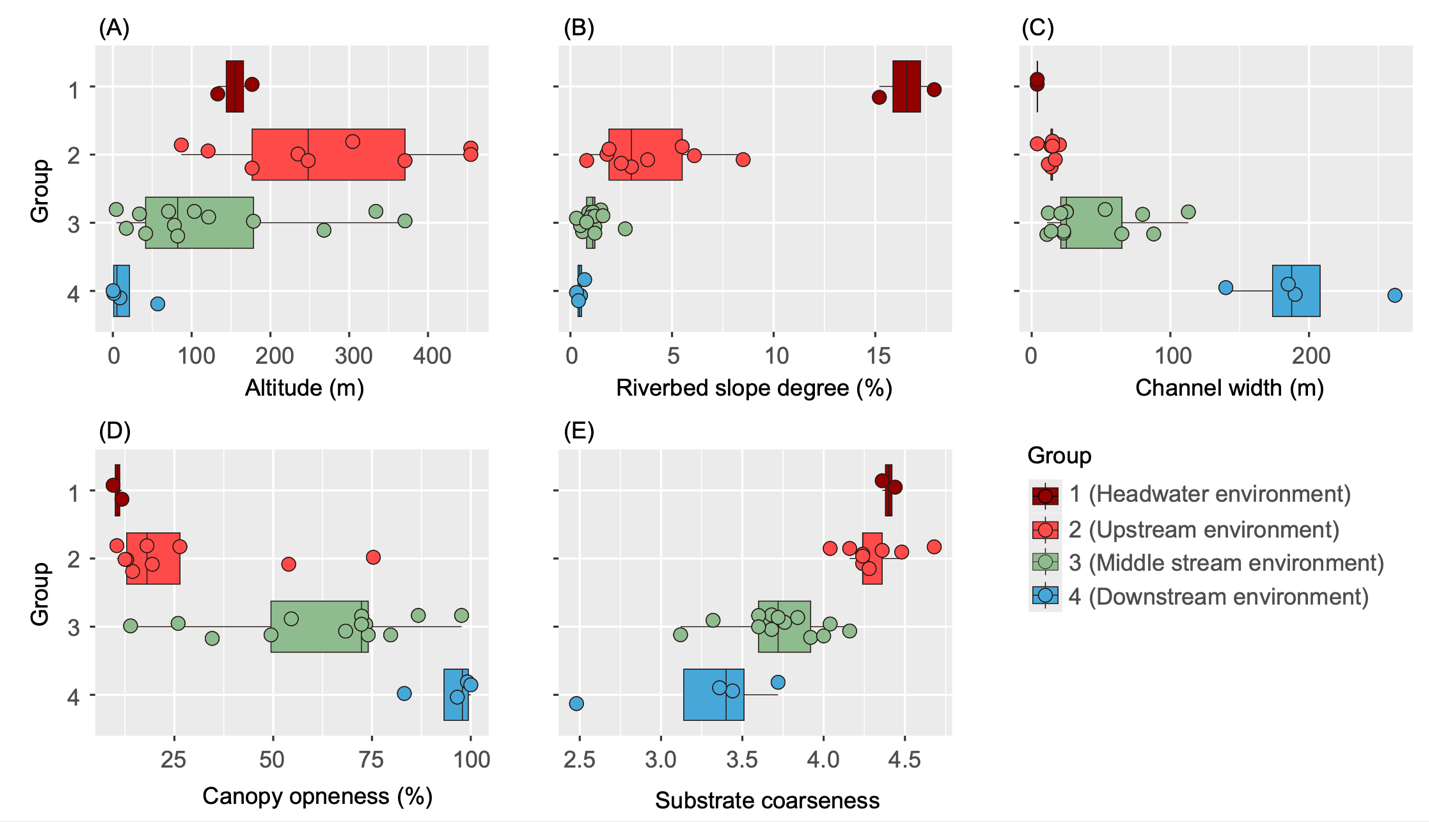

Figure S2. Box-plot showing range of environmental factors in each cluster. The range of altitude in each group (A); the range of riverbed slope degree in each group (B); the range of channel width in each group (C); the range of canopy openness in each group (D); the range of substrate coarseness in each group (E)

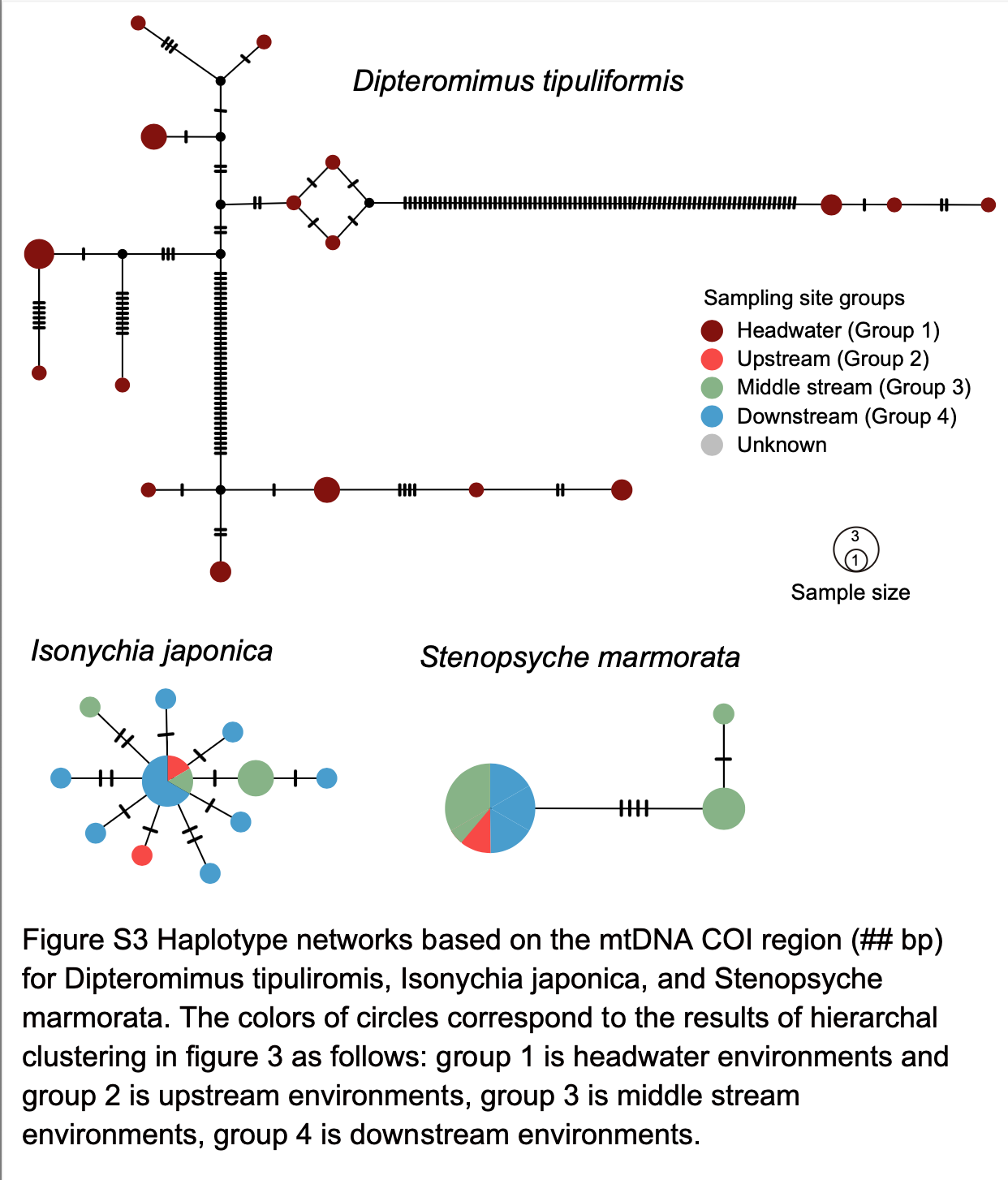
